# TORC1, BORC, ARL-8 cycling, and kinesin-1 drive vesiculation of cell corpse phagolysosomes

**DOI:** 10.1101/2022.02.09.479694

**Authors:** Gholamreza Fazeli, Roni Levin-Konigsberg, Michael C Bassik, Christian Stigloher, Ann M Wehman

## Abstract

The dynamics of phagolysosomes leading to cargo clearance are important to provide cells with metabolites and avoid auto-immune responses, but little is known about how phagolysosomes finally resolve cell corpses and cell debris. We previously discovered that polar body phagolysosomes tubulate into small vesicles to facilitate corpse clearance within 1.5 hours in *C. elegans*. Here, we show that vesiculation depends on activating TORC1 through amino acid release by the solute transporter SLC-36.1. Downstream of TORC1, BLOC-1-related complex (BORC) recruits the Arf-like GTPase ARL-8 to the phagolysosome for tubulation by kinesin-1. We find that disrupting the regulated GTP-GDP cycle of ARL-8 reduces tubulation, delays corpse clearance, and mislocalizes ARL-8 away from lysosomes. We also demonstrate that mammalian phagocytes use BORC to promote phagolysosomal degradation, confirming the conserved importance of this pathway. Finally, we show that HOPS is required for rapid degradation of the small phagolysosomal vesicles. Thus, by observing single phagolysosomes over time, we identified the molecular pathway regulating phagolysosome vesiculation that promotes efficient resolution.

## Introduction

Phagocytosis is an essential part of the innate immune response to pathogens as well as a key element for tissue homeostasis through the clearance of dying cells and debris. Billions of cell corpses are removed from the human body every day, which is important to avoid auto-immune responses (Fond and Ravichandran 2016). Many cell types, including undifferentiated cells, use phagocytosis to engulf and eventually degrade phagocytosed cargo (Fond and Ravichandran 2016; Ghose and Wehman 2021). The mechanisms of phagocytic engulfment and phagosome-lysosome fusion are well studied, but there is relatively little known about how later stages of phagolysosomal dynamics influence cargo degradation, which are important for nutrient acquisition and to modulate the immune response (Martinez et al. 2016). The final stage where a phagolysosome resolves, degrading all cargo and recycling its lipid membrane to endolysosomal pathways, is particularly mysterious due to challenges in observing these late stages in phagolysosome resolution. In most experimental systems, it is difficult to predict when a phagocytic event will occur and unambiguously track an individual phagolysosome for long periods. Furthermore, the widely used study of non-degradable phagocytic cargos, such as inorganic beads, has precluded analysis of the ultimate stages of phagocytic clearance.

We have shown that a single cell corpse, the second polar body, is stereotypically phagocytosed by large blastomeres within a 15-minute window in early *C. elegans* embryos (Fazeli et al. 2018). We found that cargo degradation occurs rapidly, ∼90 minutes after phagosome-lysosome fusion, allowing us to image phagolysosomal dynamics during resolution *in vivo*. We discovered that phagolysosomes tubulate to form small vesicles that rapidly degrade corpse proteins (Fig. 1A-B, Supplemental Movie 1) (Fazeli et al. 2018). When vesiculation was inhibited, phagolysosomal degradation was delayed by over an hour (Fazeli et al. 2018), revealing that phagolysosomal vesiculation promotes cargo degradation. This process is distinct from lysosome reformation after autophagy or phagocytosis (Yu et al. 2010; Gan et al. 2019), as the phagolysosomal vesicles contain undegraded cargo (Krajcovic et al. 2013; Fazeli et al. 2018; Lancaster et al. 2021).

**Fig. 1:**
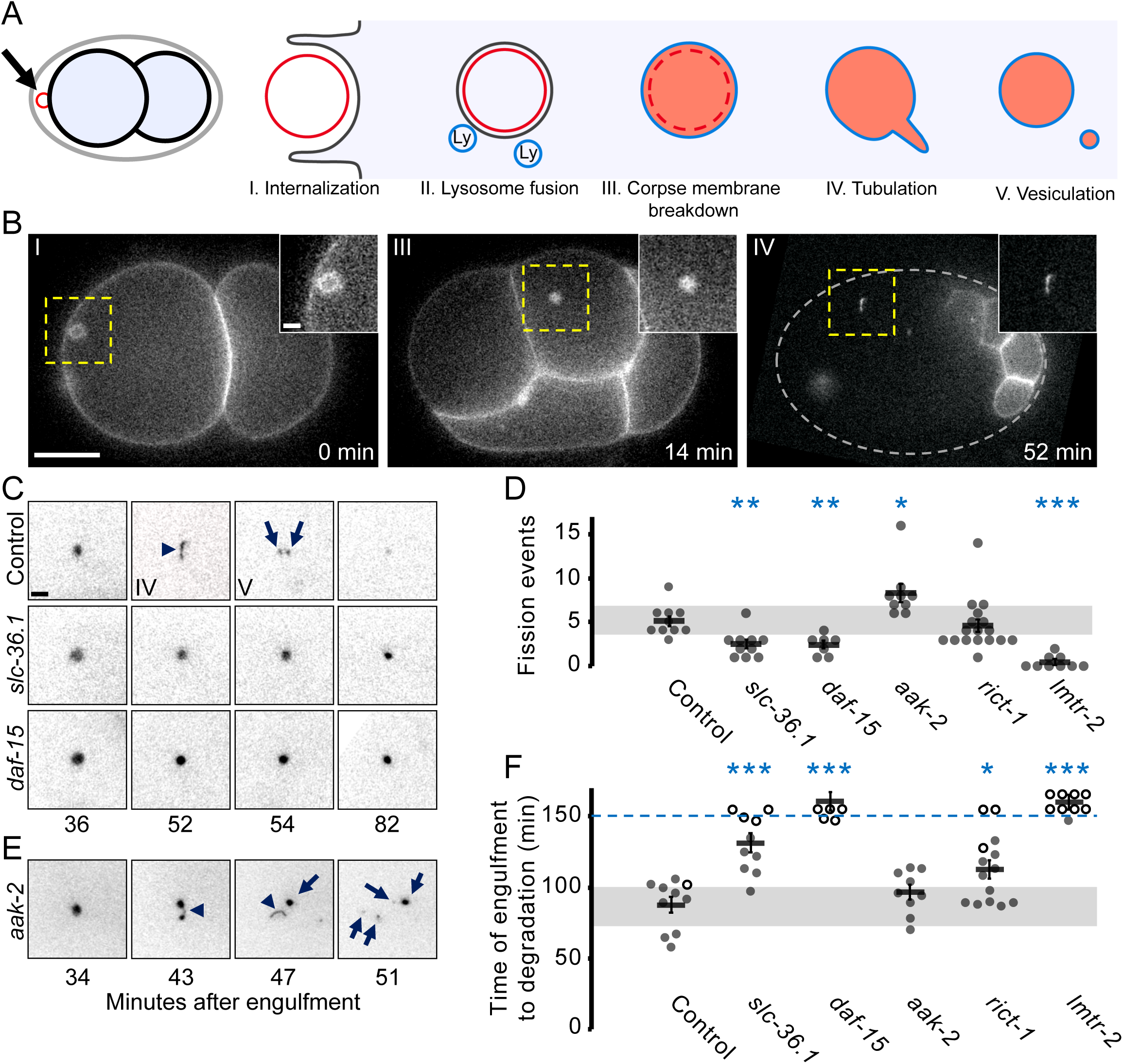
Phagolysosome vesiculation requires amino acid release and TORC1 activation. A) Schematic representation of the fate of the second polar body corpse. The corpse is phagocytosed (I) by a blastomere cell and the phagosome is fused with lysosomes (II). Next the corpse membrane (red) breaks down inside the phagolysosome membrane (blue), dispersing the membrane tag inside the lumen of the phagolysosome (III). Around 40 min later tubules containing cargo are extended from phagolysosome (IV), which then are released as small vesicles (V) to facilitate degradation. B) Representative images of an embryo expressing mCh::PH::ZF1 with a corpse (yellow square) at the stages of internalization (I), corpse membrane breakdown (III) and phagolysosome tubulation (IV). Scale bars in the main panel and magnified inset show 10 and 2 μm respectively. Indicated times are minutes after engulfment. Grey dotted line in the left panel shows the outline of the embryo. See also Supplemental Movie 1. C) The polar body phagolysosome tubulates (arrowhead) to release vesicles (arrows) in a control embryo, but not after knockdown of the lysosomal amino acid transporter SLC-36.1 or a TORC1 subunit RAPTOR/DAF-15. Panels are inverted for better contrast. D) Fission events are decreased compared to control (5±1) after knockdown of SLC- 36.1 (3±0) or DAF-15 (2±0), but not after TORC2 subunit RICTOR/RICT-1 knockdown (5±1). *aak-2(ok524)* mutants show increased vesiculation (8±1). Fission events were decreased in Ragulator *lmtr-2(tm2367)* mutants (0±0). E) Disrupting the negative regulator of TORC1, AMPK/AAK-2, results in increased fission events (arrowheads) to produce more phagolysosomal vesicles (arrows). F) The 2^nd^ polar body cargo mCh::PH::ZF1 disappears from the phagolysosome 88±6 min after internalization. Timely disappearance of the corpse cargo depends on SLC-36.1 (131±7 min), DAF-15 (>150 min) and RICT-1 (113±7 min). Phagolysosome disappearance was not altered in *aak-*2*(ok524)* mutants (96±5 min). Disappearance of the phagolysosome cargo was delayed in *lmtr-2(tm2367)* mutants (>150 min). Each circle denotes one phagolysosome. Open circles denote the last frame of a time-lapse series in which the polar body phagolysosome did not disappear. Mean ± SEM is shown. Disappearance times and means beyond 150 min after engulfment are grouped above the dashed blue line. Grey bars show standard deviation of the mean in controls. *p<0.05, **p<0.01, ***p<0.001 compared to control embryos using Student’s t-test with Bonferroni correction.

In our initial study, we found that the mTOR ortholog LET-363 was required for tubulation and vesiculation of the polar body phagolysosome (Fazeli et al. 2018), consistent with observations in cultured mouse macrophages (Krajcovic et al. 2013). Decreased vesiculation of the phagolysosome delayed degradation of its protein content (Fazeli et al. 2018), revealing the importance of subdividing large phagolysosomes to ensure efficient degradation. mTOR is found in two lysosomal complexes, mTORC1 and mTORC2, which are activated by different signals and activate distinct downstream signaling targets (Saxton and Sabatini 2017). However, it is unknown which proteins upstream or downstream of mTOR are involved in phagolysosome vesiculation.

We also discovered that the small ARF-like GTPase ARL-8 is required for phagolysosome tubulation and cargo degradation (Fazeli et al. 2018). Similar to other small GTPases, ARL-8 cycles between a GDP-bound and GTP-bound state (Khatter et al. 2015b), and GTP-bound ARL-8 can interact with kinesin motor proteins to mediate axonal transport of synaptic vesicles (Klassen et al. 2010; Niwa et al. 2016). Intriguingly, mammalian ARL8 localizes to lysosomes and promotes their anterograde transport by kinesins (Rosa-Ferreira and Munro 2011). Therefore, we hypothesized that ARL-8 links the phagolysosome membrane to motor proteins on microtubules to extend tubules and release vesicles (Fazeli et al. 2018), however it was unclear whether GTP-bound ARL-8 was sufficient to drive phagolysosomal tubulation or whether TOR and ARL-8 were part of the same pathway regulating tubulation.

During organelle transport, ARL8 has been connected to the BLOC-1-related complex BORC (Pu et al. 2015; Farias et al. 2017; Niwa et al. 2017; De Pace et al. 2020). BORC is a multi-subunit complex that localizes on the lysosomal membrane via the myristoyl group of its mammalian subunit Myrlysin (Pu et al. 2015). The Myrlysin ortholog SAM-4 binds to nucleotide free ARL-8, recruiting ARL-8 to the lysosome, and is thought to act as a guanosine exchange factor (GEF) (Niwa et al. 2017). ARL8 recruitment to lysosomes also depends on the BORC subunit Lyspersin (Filipek et al. 2017; Pu et al. 2017), which interacts with LAMTOR/Ragulator, a lysosomal complex that recruits and activates mTOR (Filipek et al. 2017; Pu et al. 2017) (reviewed in (Colaco and Jaattela 2017)). However, it was unknown whether BORC could regulate phagolysosome tubulation.

Here, we report that phagolysosome tubulation occurs after amino acid release by solute carriers, which in turn activate TORC1. Downstream of TORC1, BORC activates ARL- 8 for tubulation of the phagolysosome by kinesin-1. Both GDP- and GTP-locked mutants of *arl-8* show reduced tubulation of the phagolysosome and disrupt ARL-8 localization, revealing that ARL-8 needs to cycle to promote tubulation on phagolysosomes. We also discovered that BORC proteins are required to resolve phagolysosomes in mammalian cells. Finally, the HOPS tethering complex, an ARL-8 effector that promotes lysosome fusion (Khatter et al. 2015b), is required to rapidly degrade the small phagolysosomal vesicles released by phagolysosome tubulation. Thus, using an *in vivo* model to observe resolution of a single phagolysosome, we have identified a conserved molecular pathway regulating tubulation and vesiculation to promote degradation of phagolysosome cargos.

## Results

### Phagolysosome vesiculation requires amino acid release and TORC1 activation

To determine which TOR complex is involved in phagolysosome tubulation in *C. elegans*, we depleted TORC1 and TORC2 subunits that define the substrate specificity of each complex, specifically the Raptor homolog DAF-15 for TORC1 and the Rictor homolog RICT-1 for TORC2 and examined polar body phagolysosome dynamics. Similar to the two-fold decrease in phagolysosome fission events we saw after depleting TOR (Fazeli et al. 2018), treatment with *daf-15* RNAi resulted in a two-fold decrease in fission events (Fig. 1C-D). We also saw that phagolysosomes remained large and that the degradation of polar body protein cargo was delayed over 1 hour after depleting TOR (Fazeli et al. 2018), as well as DAF-15 (Fig. 1C, F). In contrast, *rict-1* RNAi did not significantly affect vesiculation (Fig. 1D) but did lead to a half hour delay in phagolysosomal degradation (Fig. 1F). These findings suggest that only TORC1 is required for phagolysosome vesiculation, but both TORC1 and TORC2 play important roles in phagolysosomal degradation.

To confirm that the TORC1 pathway is involved in vesiculation, we next disrupted upstream regulators of TORC1. TORC1 is activated by amino acid release by solute carriers, including SLC-36.1 (Gan et al. 2019), and negatively regulated by AMP Kinase (Kim et al. 2011). Knockdown of *slc-36.1* decreased the number of fission events by half and delayed degradation of the phagolysosome over 40 minutes (Fig. 1C-D, F), consistent with amino acid release promoting vesiculation. In contrast, deleting the AMPK catalytic subunit AAK-2 resulted in a 60% increase in vesiculation in *aak-2(ok524)* mutants (Fig. 1D-E), consistent with AAK-2 negatively regulating TORC1 in phagolysosome vesiculation. Interestingly, increased vesiculation did not lead to faster clearance of polar body contents (Fig. 1F), suggesting that other lysosomal factors are limiting for protein degradation. Thus, activation and inhibition of TORC1 activity modulates vesiculation and degradation of phagolysosomal cargos.

### BORC is required for phagolysosome tubulation and resolution

mTOR activates ARL8 via the lysosome-resident BORC complex during organelle transport (Pu et al. 2015; Niwa et al. 2017), but BORC was not known to play a role in tubulation or vesiculation. We asked whether BORC subunits play a role in phagolysosome vesiculation, starting with the Myrlysin homolog SAM-4 (Niwa et al. 2017). In *sam-4(tm3828)* mutants, phagolysosomes remained large (Fig. 2A). We observed a five-fold decrease in phagolysosome vesiculation (Fig. 2B), and degradation of the phagolysosome contents was delayed over an hour (Fig. 2C, Table S1). Loss-of-function mutants for BORC subunits *BORCS1/blos-1(ok3707)*, *BORCS2/blos-2(js1351)*, and *Lyspersin/blos-7(wy1159)* also reduced vesiculation by five-fold and delayed degradation over an hour (Fig. 2A-C, Supplemental Movie 2, Table S1). These results demonstrate that the major subunits of BORC are required for phagolysosome vesiculation and the timely degradation of a cell corpse.

**Fig. 2:**
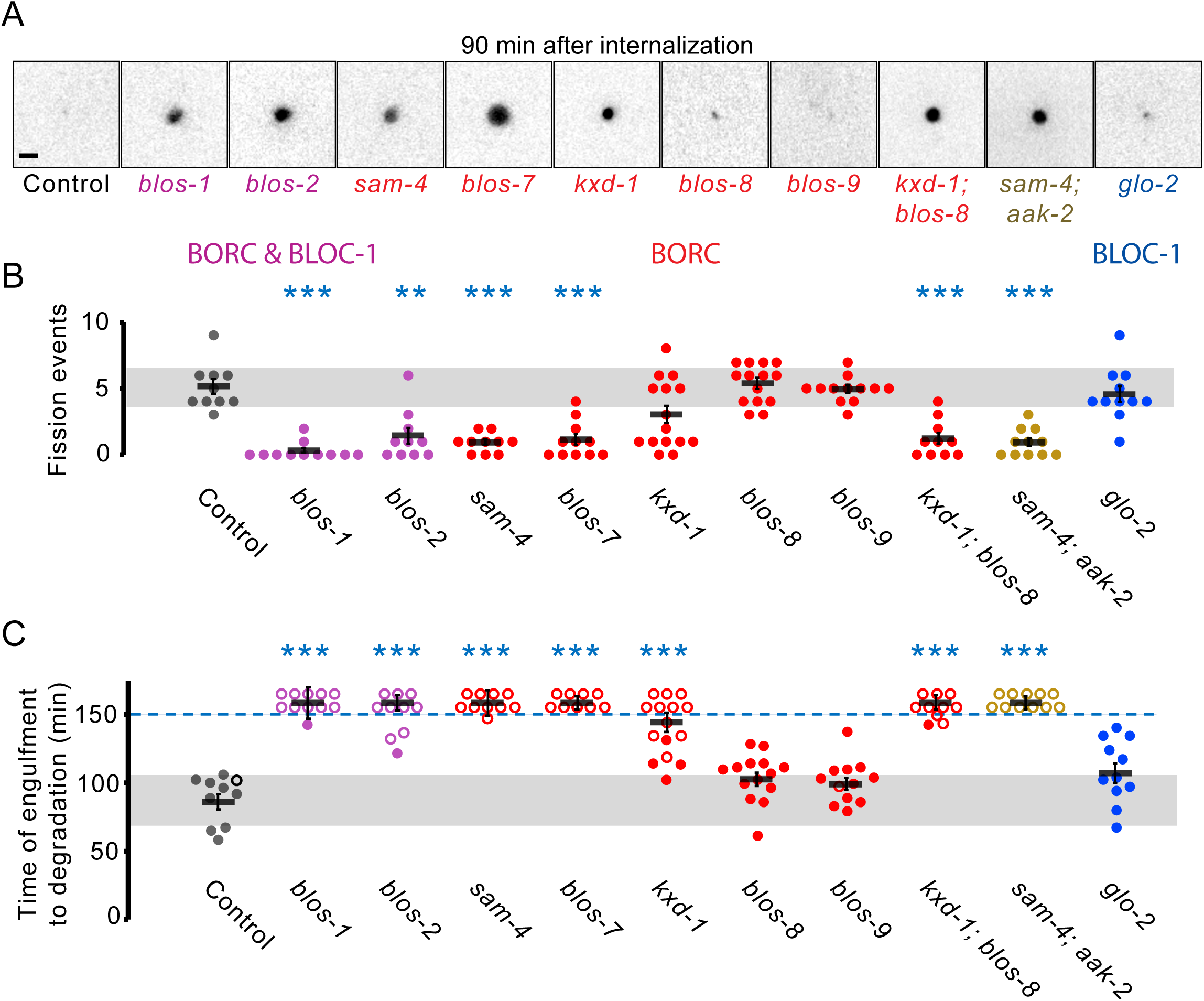
BORC but not BLOC-1 is required for phagolysosome tubulation and cargo degradation. A) Inverted images of corpse phagolysosomes 90 minutes after engulfment in control and indicated mutants. Scale bar is 2 μm. B) Phagolysosomal fission events were reduced compared to control embryos (5±1) in BORC subunit *blos-1(ok3707)* (0±0), *blos-2(js1351)* (1±1), *sam-4(tm3828)* (1±0), or *blos-7(wy1159)* (1±0) mutants. Single *kxd-1(js1356)* (3±1), *blos-8(js1354)* (5±0) or *blos-9(js1352)* (5±0) mutants did not significantly decrease fission events, but *kxd-1(js1356); blos-8(js1354)* double mutants decreased fission events (1±0), implying redundancy of some BORC subunits. Vesiculation was disrupted in *sam-4(tm3828); aak-2(ok524)* double mutants (1±0), which was significantly different from *aak-2(ok524)* single mutants (p<0.001). Disrupting the related complex BLOC-1 with *glo-2(zu455)* mutants did not affect vesiculation (5±1). C) The 2^nd^ polar body cargo mCh::PH::ZF1 disappears from the phagolysosome 88±6 min after internalization. BORC mutants *blos-1(ok3707)*, *blos-2(js1351)*, *sam-4(tm3828)*, *blos-7(wy1159)* (all >150 min), or *kxd-1(js1356)* (146±7 min), delayed cargo disappearance but not *blos-8(js1354)* (104±5 min) or *blos-9(js1352)* (101±5 min) mutants. Cargo degradation was delayed >150 min in *kxd-1(js1356); blos-8(js1354)* or *sam-4(tm3828); aak-2(ok524)* double mutants. Disappearance of the phagolysosome cargo was not affected in *glo-2(zu455)* mutants (108±7 min). Open circles denote the last frame of a time-lapse series in which the polar body phagolysosome did not disappear. Mean ± SEM is shown. Disappearance times and means beyond 150 min after engulfment are grouped above the dashed blue line. Grey bars show standard deviation of the mean in controls. **p<0.01, ***p<0.001 compared to control embryos using Student’s t-test with Bonferroni correction. See also Supplemental Movie 2.

BORC also contains several small subunits that have disparate roles in synaptic vesicle or lysosome transport: only KXD1 and Diaskedin were required for lysosome transport (Pu et al. 2015), while only the MEF2BNB ortholog BLOS-9 was required for synaptic vesicle transport (Niwa et al. 2017). We found that phagolysosome tubulation and degradation occurred normally in mutants for the Diaskedin homolog *blos-8(js1354)* and for *blos-9(js1352)* (Fig. 2A-C, Table S1), while degradation of the phagolysosome was delayed almost an hour in *kxd-1(js1356)* mutants (Fig. 2C, Table S1). These findings suggest that the small BORC subunits have separable functions in synaptic vesicle transport, lysosome transport, or phagolysosomal degradation. Interestingly, *kxd-1(js1356)* mutants showed a bimodal vesiculation phenotype, with half normal and half disrupted (Fig. 2B), leaving open the possibility that the small subunits also play redundant roles in phagolysosomal vesiculation. To test this, we generated *blos-8(js1354); kxd-1(js1356)* double mutant embryos and discovered that phagolysosome vesiculation was reduced five-fold and cargo degradation was delayed over an hour, similar to the major BORC subunits (Fig. 2A-C, Table S1). These data indicate that the small BORC subunits have distinct, but partially overlapping functions and implicate the subunits involved in lysosome transport in phagolysosome degradation.

BLOS-1 and BLOS-2 also act in a related complex BLOC-1, which is involved in the biogenesis of lysosome-related organelles (Langemeyer and Ungermann 2015). Therefore, we tested whether the BLOC-1-specific subunit Pallidin/GLO-2 has a role in phagolysosome tubulation. Vesiculation and degradation occurred normally in strong loss-of-function *glo- 2(zu455)* mutants (Fig. 2A-C, Table S1), suggesting that BLOS-1 and BLOS-2 act as part of the BORC complex in phagolysosome resolution.

The BORC subunit Lyspersin has been shown to bind to LAMTOR2 (Filipek et al. 2017; Pu et al. 2017), part of the Ragulator complex that acts as a GEF for Rag GTPases to recruit mTORC1 to lysosomes (Sancak et al. 2010; Bar-Peled et al. 2012). LAMTOR2 binding to Lyspersin inhibits BORC to prevent ARL8-mediated lysosome transport (Filipek et al. 2017; Pu et al. 2017). Therefore, we asked whether disrupting LMTR-2 would activate BORC and bypass the requirement for TORC1 in phagolysosome vesiculation. In *lmtr-2*(*tm2367*) deletion mutants, phagolysosome vesiculation was reduced over five-fold and degradation was delayed by over an hour (Fig. 1D, F), suggesting that releasing LMTR-2 inhibition of BLOS-7 was not sufficient to allow BORC and ARL-8 to promote phagolysosome resolution.

To test whether BORC acts downstream of the TORC1 pathway in phagolysosome resolution, we asked whether deleting *sam-4* could suppress TORC1 activation caused by disrupting AMPK. The polar body phagolysosome in *sam-4(tm3828); aak-2(ok524)* double mutants showed a five-fold reduction in vesiculation and degradation delayed by over an hour (Fig. 2A-C), which was significantly different from controls or *aak-2* single mutants (p<0.001). These data place BORC downstream of TORC1 activation in regulating phagolysosome vesiculation.

### The ARL-8 GTPase must cycle between GDP- and GTP-bound states for phagolysosome tubulation

We next asked whether ARL-8 acts downstream of TORC1 in phagolysosome tubulation using a partial loss-of-function *arl-8(wy271)* mutant lacking 0.2 kb of the promoter, the 5’ UTR, and the first two codons of *arl-8* (Fig. 3A) (Klassen et al. 2010). Consistent with *arl-8* knockdown (Fazeli et al. 2018), phagolysosome vesiculation was reduced five-fold and degradation was delayed over an hour in *arl-8(wy271)* deletion mutants (Fig. 3B-D). The *arl- 8(wy271); aak-2(ok524)* double mutants showed similarly decreased phagolysosome fission events and delayed degradation (Fig. 3B-D), confirming that both ARL-8 and BORC act downstream of TORC1 activation in phagolysosome resolution.

**Fig. 3:**
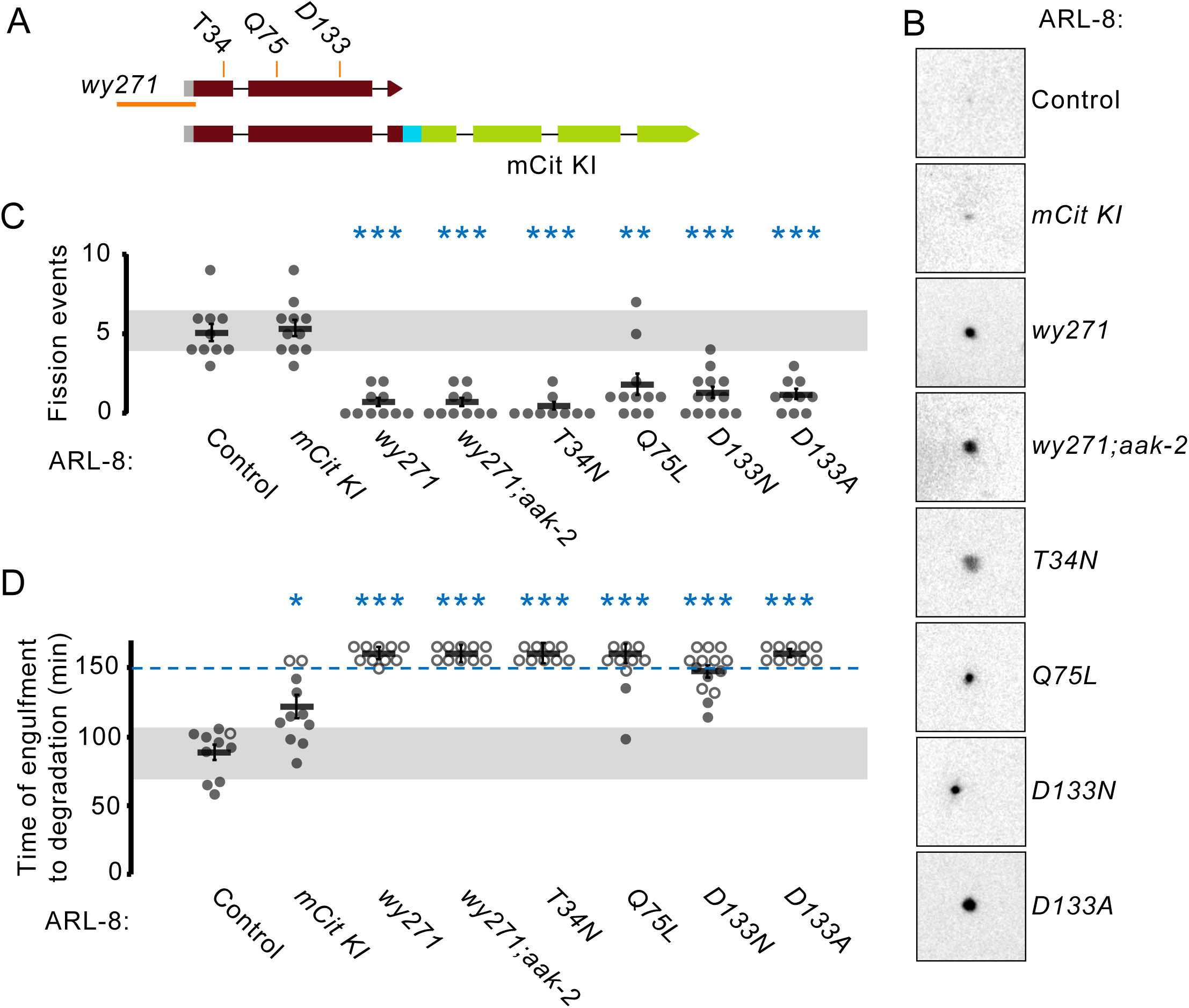
Dysregulating the ARL-8 GTPase cycle disrupts phagolysosome tubulation. A) Genomic *arl-8* locus with *wy271* deletion (orange line) and the point mutations T34N (GDP-bound), Q75L (GTP-bound), D133N and D133A mutations (G4 motif) indicated. mCitrine was inserted in the C-terminus of *arl-8* with a flexible linker (cyan) with or without the above point mutations. B) Inverted images of corpse phagolysosomes 90 minutes after engulfment in control and indicated mutants. Scale bar is 2 μm. C) Knocking mCit into *arl-8* did not affect phagolysosomal fission events (5±1 vs. 5±1 in controls), while fission events were dramatically reduced in *arl-*8(*wy271)* (1±0), *T34N* (0±0), *Q75L* (2±1), *D133N* (1±0) or *D133A* (1±0) mutants. Vesiculation was also disrupted in *arl-8(wy271); aak-2(ok524)* double mutants (1±0), which was significantly different from *aak-2(ok524)* mutants (p<0.001). D) The ARL-8::mCit knock-in slightly delayed the disappearance of the 2^nd^ polar body phagolysosome cargo marker mCh::PH::ZF1 (120±8 min), compared to control (88±6 min), whereas cargo degradation was severely delayed in *arl-*8(*wy271)*, *T34N*, *Q75L*, *D133A* (all >150 min) or *D133N* (146±4) mutants, as well as *arl-8*(*wy271); aak-2(ok524)* double mutants (>150 min). Open circles denote the last frame of a time-lapse series in which the polar body phagolysosome did not disappear. Mean ± SEM is shown. Disappearance times and means beyond 150 min after engulfment are grouped above the dashed blue line. Grey bars show standard deviation of the mean in controls. *p<0.05, **p<0.01, ***p<0.001 compared to control embryos using Student’s t-test with Bonferroni correction.

As BORC recruits nucleotide-free ARL-8 and is thought to promote GTP exchange for ARL-8 (Niwa et al. 2017), we asked how the state of the bound nucleotide dictates the activity of ARL-8 during phagolysosome tubulation. We first generated point mutations in ARL-8 which are predicted to lock the GTPase in either a GDP- or GTP-bound state (Bagshaw et al. 2006). The GDP-bound *arl-8(T34N)* mutant reduced vesiculation five-fold and delayed degradation by over an hour (Fig. 3B-D), consistent with previous reports predicting it to be inactive (Bagshaw et al. 2006). Interestingly, the putative GTP-locked *arl-8(Q75L)* mutant caused a two-fold decrease in vesiculation and also delayed phagolysosome degradation over an hour (Fig. 3B-D), in contrast to previous reports that this allele can rescue synaptic vesicle trafficking defects (Klassen et al. 2010) or embryonic lethality of the strong loss-of-function *arl-8(tm2504)* deletion allele (Nakae et al. 2010). We next examined mutations in the GTPase G4 motif at Aspartate 133, which are predicted to allow nucleotide exchange independent of a GEF (Morris et al. 2018) and have been shown to rescue synaptic vesicle trafficking defects (Niwa et al. 2017). We also observed a five-fold reduction in phagolysosome vesiculation and an hour delay in cargo degradation with *arl-8(D133A)* and *arl-8(D133N)* mutants (Fig. 3B-D). These data indicate that both GDP- and GTP-locked as well as unlocked alleles disrupt phagolysosome tubulation and suggest that ARL-8 needs to cycle between GDP- and GTP-bound in a controlled fashion to regulate phagolysosome budding.

### ARL-8 localization depends on its nucleotide state

To understand how ARL-8 localization is affected by its nucleotide state, we first tagged endogenous ARL-8 with mCitrine (Fig. 3A). The ARL-8::mCit knock-in did not alter vesiculation events (Fig. 3C), suggesting that ARL-8::mCit promoted phagolysosome vesiculation normally. However, degradation of polar body phagolysosome content was delayed by half an hour (Fig. 3B, D), suggesting that the fluorescent tag may have a minor effect on another aspect of ARL- 8’s function in phagolysosomal protein degradation. ARL-8::mCit localized to punctate structures in early embryos (Fig. 4B), which we confirmed were 50 ± 6% lysosomal by colocalization with an mScarlet-I-tagged lysosome-resident BORC subunit SAM-4::mSc (Fig. 4L, S2F-G). 10 ± 1% of ARL-8::mCit also colocalized with the early endosomal small GTPase mCh::RAB-5 (Fig. 4M, Pearson coefficient 0.0003 ± 0.01), consistent with overexpression studies showing that ARL-8::GFP mainly localized to lysosomes but was also observed on other structures (Sasaki et al. 2013). ARL-8::mCit-positive vesicles started appearing on the polar body phagosome shortly after engulfment and ARL-8 appeared as a ring around the phagolysosome 6 ± 2 minutes after its engulfment (n=8, Fig. 4A), similar to the timing of LMP- 1-positive lysosome recruitment (Fazeli et al. 2018), which suggests that ARL-8 is recruited to the phagosome at lysosome fusion.

**Fig. 4:**
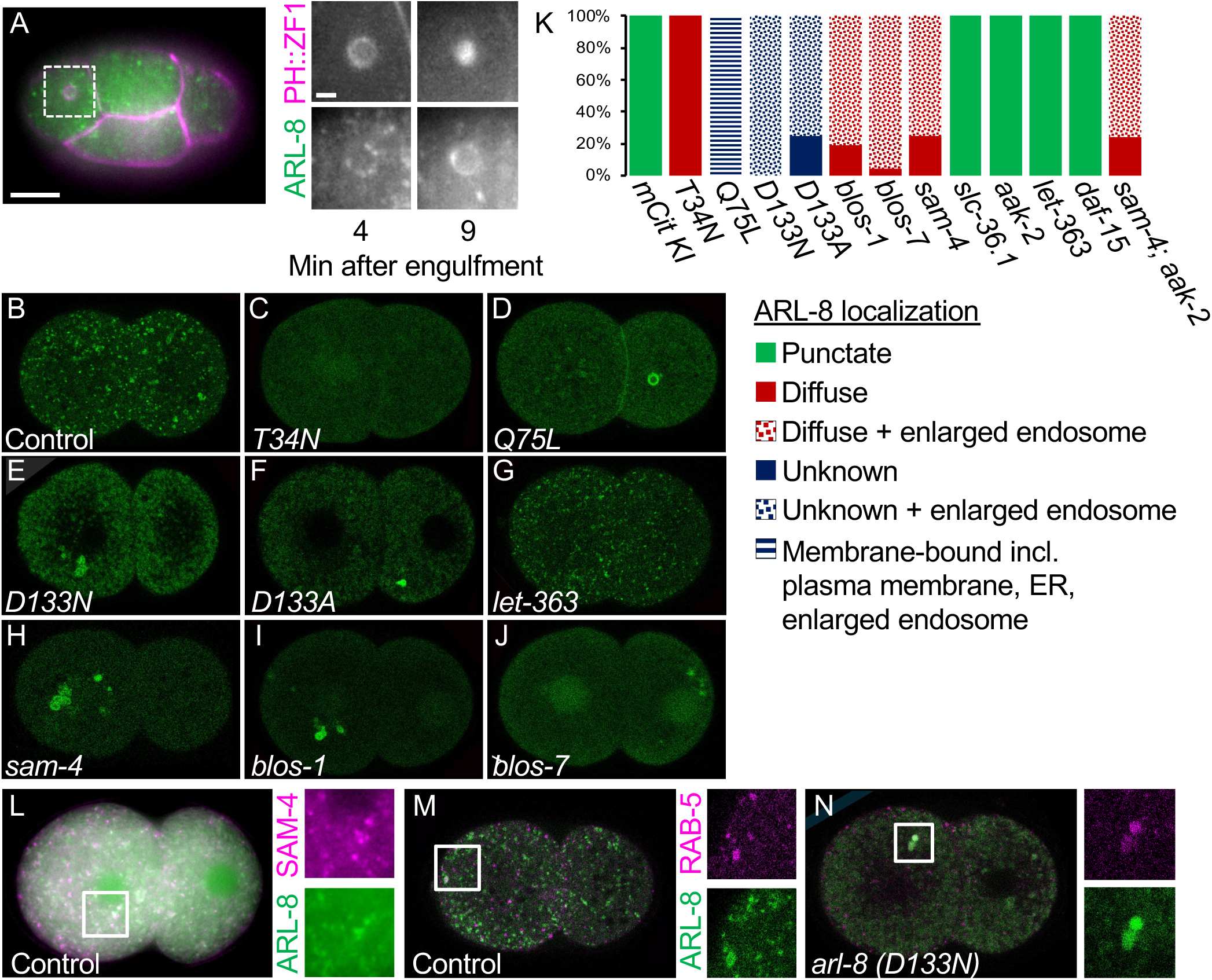
ARL-8 localization to lysosomes, phagolysosomes, and early endosomes depends on GTP cycling. A) ARL-8::mCit puncta (green) appear around the polar body membrane (mCh::PH::ZF1, magenta) shortly after internalization. By the time of the corpse membrane breakdown inside the phagolysosome, ARL-8::mCit appears as a ring around the phagosome. Scale bar is 10 μm. Insets show the magnified dashed white square at the indicated time after engulfment. Scale bar is 2 μm. B) In control embryos, endogenously tagged ARL-8::mCit localizes to puncta (lysosomes and endosomes). C) ARL-8(T34N)::mCit is mostly diffuse. D) ARL-8(Q75L)::mCit localizes to the plasma membrane, ER, and large vesicles. E-F) ARL-8(D133N)::mCit and ARL-8(D133A)::mCit associate with unknown structures in cytoplasm and large interconnected endosomes. G) ARL-8::mCit localizes to puncta after RNAi-mediated *TOR/let-*363 knockdown, similar to control. H-J) In BORC subunit *sam-4 (tm3828)*, *blos-1(ok3707)*, or *blos-7(wy1159)* mutants, ARL-8::mCit appears diffuse but also associates with large interconnected endosomes. K) Quantification of ARL-8::mCit localization in ARL-8 or BORC mutants or embryos deficient in TOR-related proteins. See also Supplemental Fig. S1 and S2. L) ARL- 8::mCit colocalizes with the lysosomal resident BORC subunit SAM-4. M-N) ARL-8::mCit infrequently colocalizes with the early endosomal marker mCh::RAB-5 (M), while the large interconnected endosomes in *arl-8(D133N)* mutants are RAB-5-positive (N).

We next generated *arl-8* point mutations in the mCit knock-in strain and examined their localization. GDP-bound ARL-8(T34N)::mCit appeared largely dispersed (Fig. 4C), consistent with the known role of GTP binding in promoting membrane association of GTPases (Cherfils and Zeghouf 2013). GTP-locked ARL-8(Q75L)::mCit localized on enlarged tubulovesicular structures, the plasma membrane and the ER (Fig. 4D, Fig. S1A), but was not observed on discrete puncta indicative of lysosomes. In contrast, unlocked ARL-8(D133N)::mCit and ARL- 8(D133A)::mCit localized to large tubulovesicular structures and other unidentified structures (Fig. 4E-F), but not the plasma membrane or ER (Fig. 4E-F, S1B). The large tubulovesicular structures colocalized with mCh::RAB-5 in *arl-8(D133N)* mutants (Fig. 4N), revealing that interfering with the regulated GDP/GTP cycle traps ARL-8 on early endosomes and alters endosomal morphology. These findings show that ARL-8 can associate to different membranes depending on its nucleotide state (Fig. 4K). Thus, a regulated cycle between the GDP- and GTP-bound states is required for ARL-8 lysosome localization and phagolysosome tubulation.

### BORC but not TORC1 is required for ARL-8 localization to lysosomes and the phagolysosome

As Myrlysin is required for ARL8 localization to lysosomes in HeLa cells (Pu et al. 2015), we asked whether SAM-4 and BORC are required for ARL-8 localization to the polar body phagolysosome. ARL-8::mCit was not observed on the polar body phagolysosome in *sam-4(tm3828)* mutants (Fig. 5A). Instead, ARL-8::mCit associated with large tubulovesicular networks in *sam-4(tm3828)* mutants (Fig. 4H), which we confirmed to be early endosomes by colocalization with mCh::RAB-5 (Fig. 5B) and not lysosomes by LMP-1 staining (Fig. 5C). ARL- 8::mCit similarly associated with enlarged tubulovesicular networks in *blos-1(ok3707)* and *blos-7(wy1159)* mutants (Fig. 4I-J). Thus, BORC is required for both the lysosomal and phagolysosomal localization of ARL-8, suggesting that the phagolysosome resolution defects in BORC mutants are likely to be caused by altering ARL-8 localization.

**Fig. 5:**
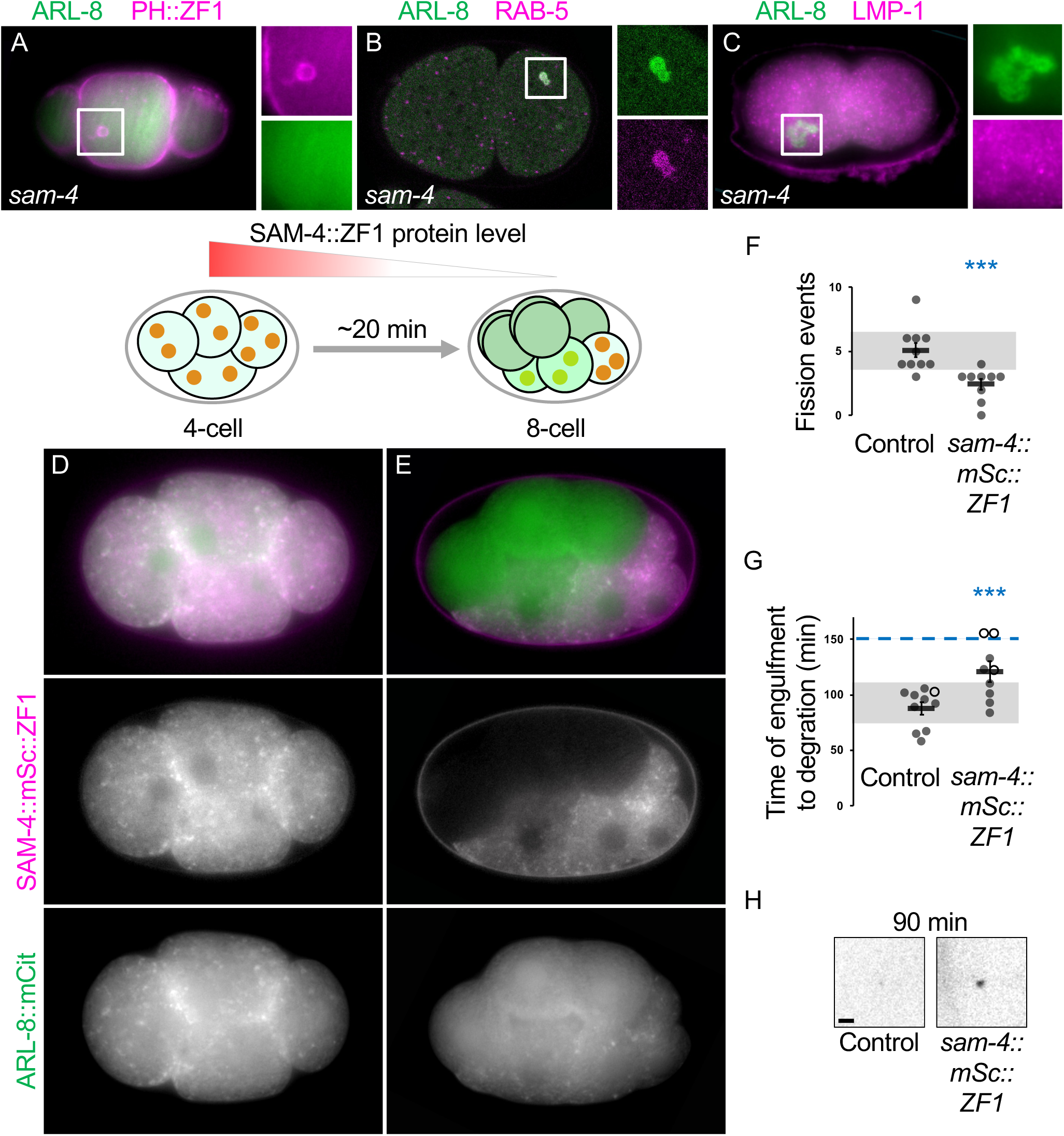
ARL-8 localization to lysosomes requires SAM-4. A) ARL-8::mCit (green) does not appear around the polar body membrane (mCh::PH::ZF1, magenta) after engulfment in *sam-4(tm3828)* mutants (see also Fig 4.A). B-C) ARL-8::mCit colocalizes with mCh::RAB-5 on large interconnected vesicles in *sam-4 (tm3828)* mutants (B), but not with lysosomal marker LMP-1 (C). D) SAM-4::mSc::ZF1 (magenta) and ARL-8::mCit (green) colocalize in puncta in a 4-cell embryo prior to onset of ZIF-1-mediated degradation. E) SAM-4::mSc::ZF1 degrades in the anterior cells of an 8-cell embryo and ARL-8::mCit disperses in the cytosol. In posterior germline cells, ZF1-tagged proteins are not degraded and the punctate pattern of SAM-4 and ARL-8::mCit and their colocalization is conserved. F) Degrading SAM-4::mSc::ZF1 significantly decreased phagolysosome fission events (2±0 vs. 5±1 in control). G) Disappearance of the 2^nd^ polar body phagolysosome cargo marker mCh::PH::ZF1 was delayed (121±9), compared to control (88±6 min). Open circles denote the last frame of a time-lapse series in which the polar body phagolysosome did not disappear. Mean ± SEM is shown. Disappearance times beyond 150 min after engulfment are grouped above the dashed blue line. Grey bars show standard deviation of the mean in controls. ***p<0.001 compared to control embryos using Student’s t-test. H) Inverted images of corpse phagolysosomes 90 minutes after engulfment in control and a SAM-4::mSc::ZF1 knock-in embryo. Scale bar is 2 μm.

Next, we asked whether TORC1 activity also alters ARL-8 localization. We saw no gross changes to ARL-8::mCit localization after disrupting TORC1 with *TOR/let-363* or *daf-15* RNAi (Fig. 4G, S2B), disrupting TORC1 activation by amino acids with *slc-36.1* RNAi (Fig. S2A), over-activating TORC1 in *aak-2(ok524)* deletion mutants (Fig. S2C), or disrupting TORC1 recruitment and BORC inhibition in *lmtr-2(tm2367)* mutants (Fig. S2D). We further confirmed that *TOR/let-363* RNAi did not significantly alter ARL-8::mCit colocalization with SAM-4 (Fig. S2E-G). These data indicate that BORC regulates ARL-8 localization independent of TORC1, leaving the molecular character of TORC1 regulation on ARL-8 unclear.

As BORC deletion would lead to chronic disruption of ARL-8 localization, which could also alter lysosomes, we used a degron approach to determine the effect of acute loss of SAM- 4 on ARL-8 localization and phagolysosomal degradation. We tagged endogenous SAM-4 with mScarlet-I and a ZF1 degron to induce ZIF-1-mediated degradation starting in anterior cells at the late 4-cell stage (Fig. 5D-E) (Armenti et al. 2014). Prior to the onset of degradation, ARL- 8::mCit localized on discrete puncta (half together with SAM-4::mSc::ZF1) (Fig. 5D). After SAM-4::mSc::ZF1 degradation, ARL-8::mCit quickly dispersed (Fig. 5E). Next, we asked whether phagolysosomal degradation is also affected by acute loss of SAM-4 and mislocalization of ARL-8. SAM-4::mSc::ZF1 embryos showed a two-fold decrease in phagolysosome fission events and delayed degradation by a half hour (Fig. 5F-H). These data indicate that SAM-4 needs to constantly promote ARL-8 localization to lysosomal and/or phagolysosomal membranes to efficiently degrade phagolysosome protein content.

### Phagolysosome vesiculation via BORC is conserved in mammals

We next asked whether proteins required for phagolysosome vesiculation in *C. elegans* are needed to vesiculate phagolysosomes in mammals. We fed murine macrophage J774 cells with opsonized sheep red blood cells (SRBC) and found that engulfed SRBC were observed 6 hours later as fragments distributed throughout the cytoplasm of engulfing cells (Fig. 6A), confirming previous work (Levin-Konigsberg et al. 2019). Next, we used CRISPR/Cas9 to mutate BORC subunits Lyspersin and Myrlysin (Supplemental Fig. 4) and found that SRBC fragmentation (stage III) was reduced in mutant clones, with more SRBCs remaining intact (stage I) or forming only a few fragments (stage II) (Fig. 6B-D). These data suggest that the role of BORC in phagolysosome vesiculation is conserved from nematodes to mammals.

**Fig. 6:**
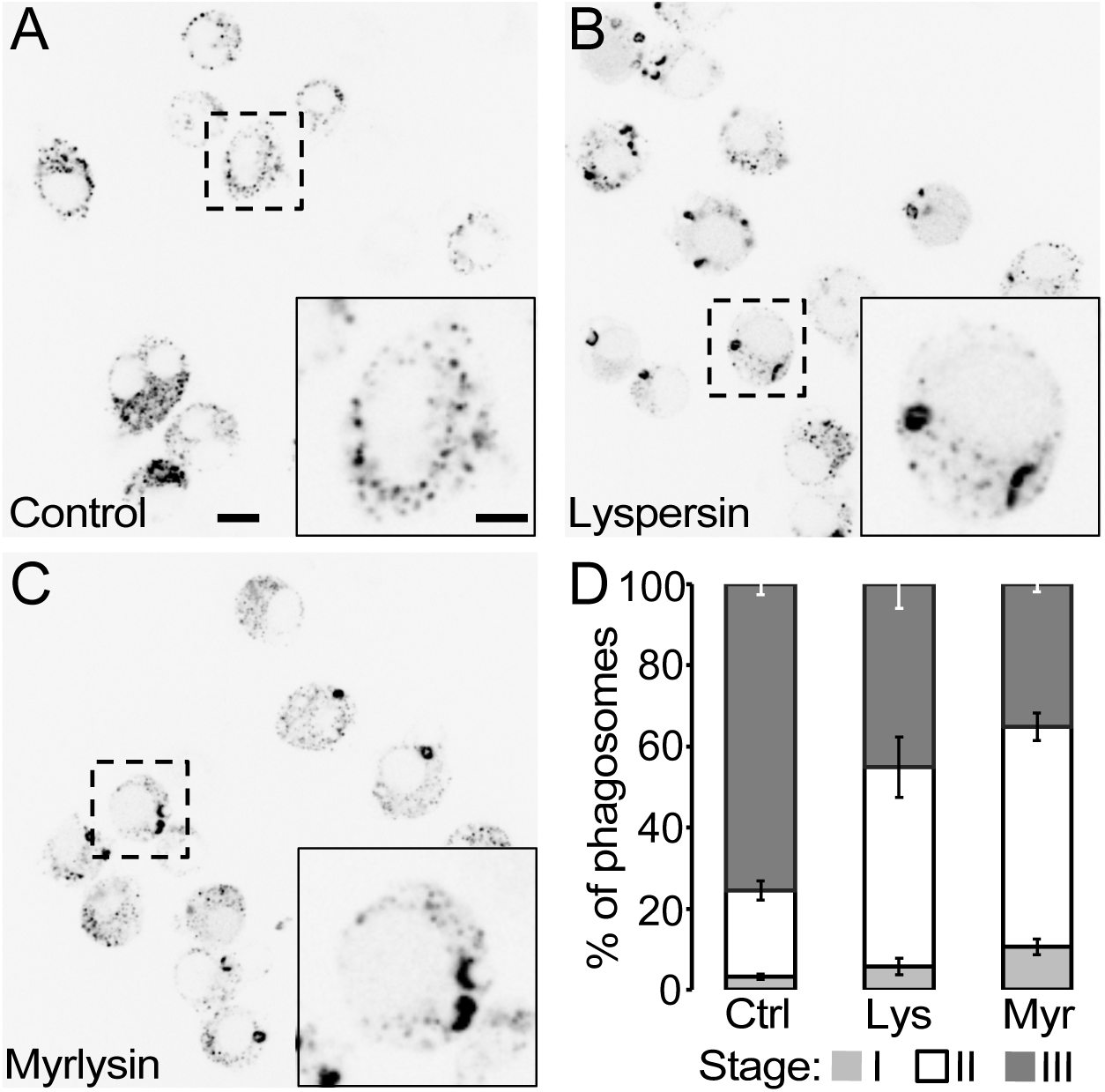
Phagolysosome vesiculation via BORC is conserved in mammals. A) Murine macrophage J774 cells treated with a control sgRNA engulfed and fragmented opsonized sheep red blood cells (SRBC) within 6 hours after feeding. B-C) Engulfed SRBC remained large with reduced fragmentation in Lyspersin (B) or Myrlysin (C) CRISPR mutant J774 cell lines. See Supplementary Fig. 4 for CRISPR edits. Dotted rectangle is magnified in the insets. Scale bars in the main image and inset are 10 and 5 μm respectively. D) Quantification of phagosome resolution revealed that Lyspersin and Myrlysin mutants significantly decreased stage III (mostly fragmented SRBC, dispersed throughout the cytoplasm) and increased stage I (large, seemingly undigested SRBC) & stage II (large partially digested SRBC with proximal small, fragmented structures). Chi-squared test, p<0.00001. Mean ± SD is shown.

### Kinesin-1 is required for phagolysosome tubulation

Finally, we asked which proteins drive phagolysosome tubulation downstream of ARL-8. Mammalian ARL8 binds to SKIP/PLEKHM2 for anterograde transport by kinesin-1 (Rosa-Ferreira and Munro 2011), as well as binding kinesins directly (Niwa et al. 2016). We screened kinesin-1 using the KIF5 ortholog UNC-116 and kinesin-3 using the KIF1 ortholog UNC-104. As strong loss-of-function *unc-116* mutants disrupt the birth of the polar body during meiosis (Yang et al. 2005), we generated an UNC-116::mCit::ZF1 strain, which degrades UNC-116 after the polar body is engulfed (Fig. 7A). Phagolysosome vesiculation showed a 2-fold decrease after UNC-116::mCit::ZF1 degradation and the resolution of phagolysosomal cargo was delayed by 30 minutes (Fig. 7B-D). In contrast, neither the number of fission events nor the timing of degradation was affected in the *unc-104(e1265)* reference allele (Fig. 7B-C). These data demonstrate that kinesin-1, not kinesin-3, is required for phagolysosome vesiculation.

**Fig. 7:**
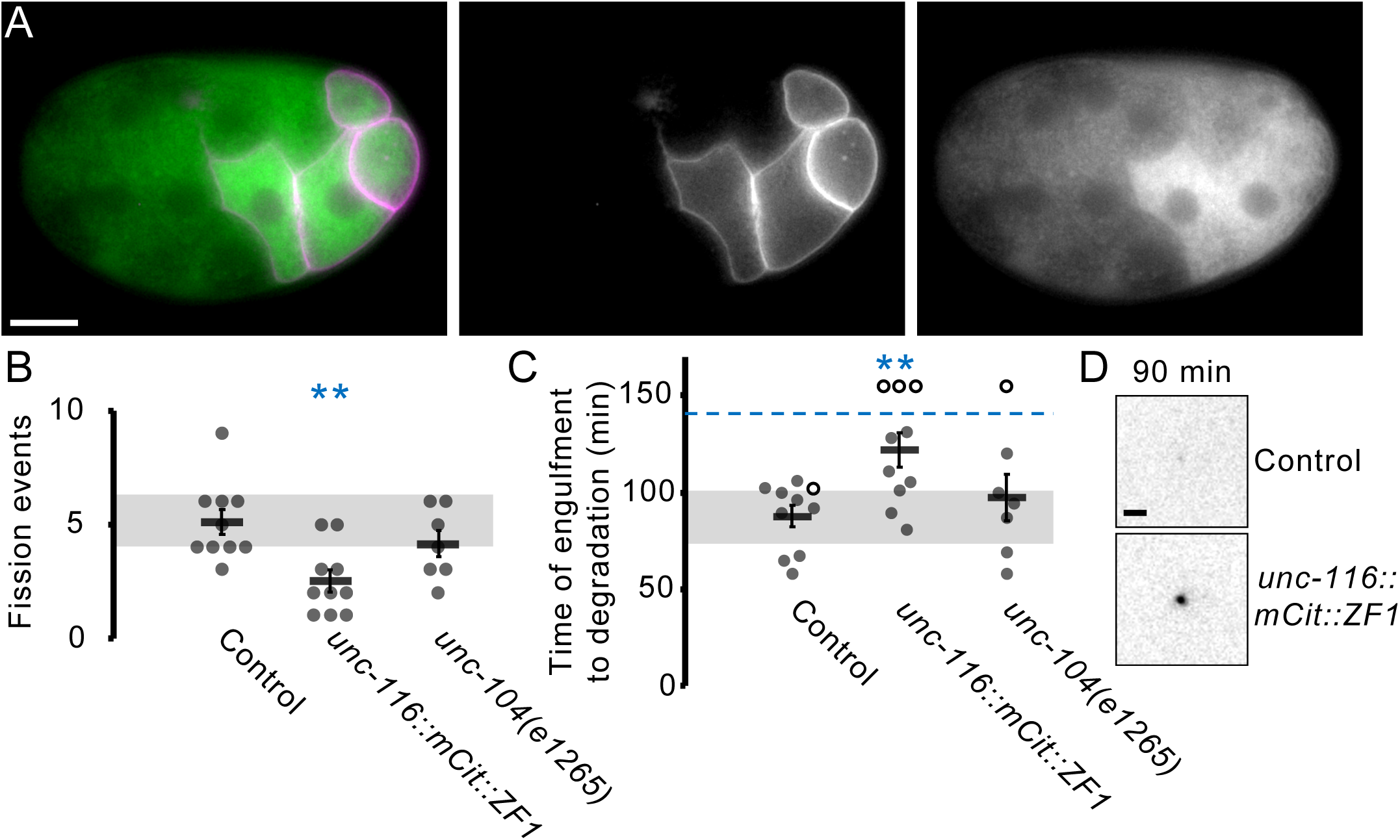
Kinesin-1 drives phagolysosome vesiculation and degradation. A) The KLC5 ortholog UNC-116::mCit::ZF1 (green) degrades in the anterior cells of a 16-cell embryo. mCh::PH::ZF1 (magenta) marks the membrane of posterior cells of the embryo, where UNC-116::mCit::ZF1 is not degraded. B) Degrading UNC-116::mCit::ZF1 significantly decreased phagolysosome fission events (3±0), whereas KLC1 mutants *unc-14(e1265)* did not differ from controls (4±1 vs. 5±1 in control). C) Disappearance of the 2^nd^ polar body phagolysosome cargo marker mCh::PH::ZF1 was delayed (122±9 min) in UNC- 116::mCit::ZF1, but not in *unc-14(e1265)* (97±12 min), compared to control (88±6 min). Open circles denote the last frame of a time-lapse series in which the polar body phagolysosome did not disappear. Mean ± SEM is shown. Disappearance times beyond 150 min after engulfment are grouped above the dashed blue line. Grey bars show standard deviation of the mean in controls. **p<0.01 compared to control embryos using Student’s t-test with Bonferroni correction. D) Inverted images of corpse phagolysosomes 90 minutes after engulfment in control and *unc-116::mCit::ZF1* embryos. Scale bar is 2 μm.

We next examined two PLEKHM family proteins, CUP-14 and RUB-1, for their role in phagolysosome resolution. Phagolysosome vesiculation was not decreased and resolution was not significantly delayed in *cup-14(cd32)* mutants or after *rub-1* RNAi treatment (Supplemental Fig. 3A-B). Knocking down *rub-1* in *cup-14* mutants also did not decrease phagolysosome vesiculation or delay phagolysosomal resolution (Supplemental Fig. 3A-B). These data suggest that PLEKHM proteins are not ARL-8 effectors during phagolysosome vesiculation in *C. elegans*.

### The ARL-8 effector HOPS is required for rapid degradation of phagolysosomal vesicles

As the vesicle-tethering HOPS complex is another effector of mammalian ARL8, with VPS41 binding ARL8B (Khatter et al. 2015a), we asked whether VPS-41 is required for phagolysosome vesiculation. RNAi knockdown of *vps-41* did not affect phagolysosome vesiculation (Fig. 8A) but significantly delayed the degradation of polar body cargo (Fig. 8B- C). Furthermore, *vps-41* knockdown specifically delayed the disappearance of phagolysosomal vesicles (Fig. 8D). Given that HOPS is best known for its role in lysosome fusion (Nguyen and Yates 2021), HOPS depletion likely affects fusion of small phagolysosomal vesicles to lysosomes. However, *vps-41* knockdown had no effect on polar body membrane breakdown within the large phagolysosome (Fig. 8E), an earlier process that depends on RAB- 7-mediated phagosome-lysosome fusion (Fazeli et al. 2018). Together, these data suggest that the requirement for HOPS during lysosome fusion may depend on the target vesicle.

**Fig. 8:**
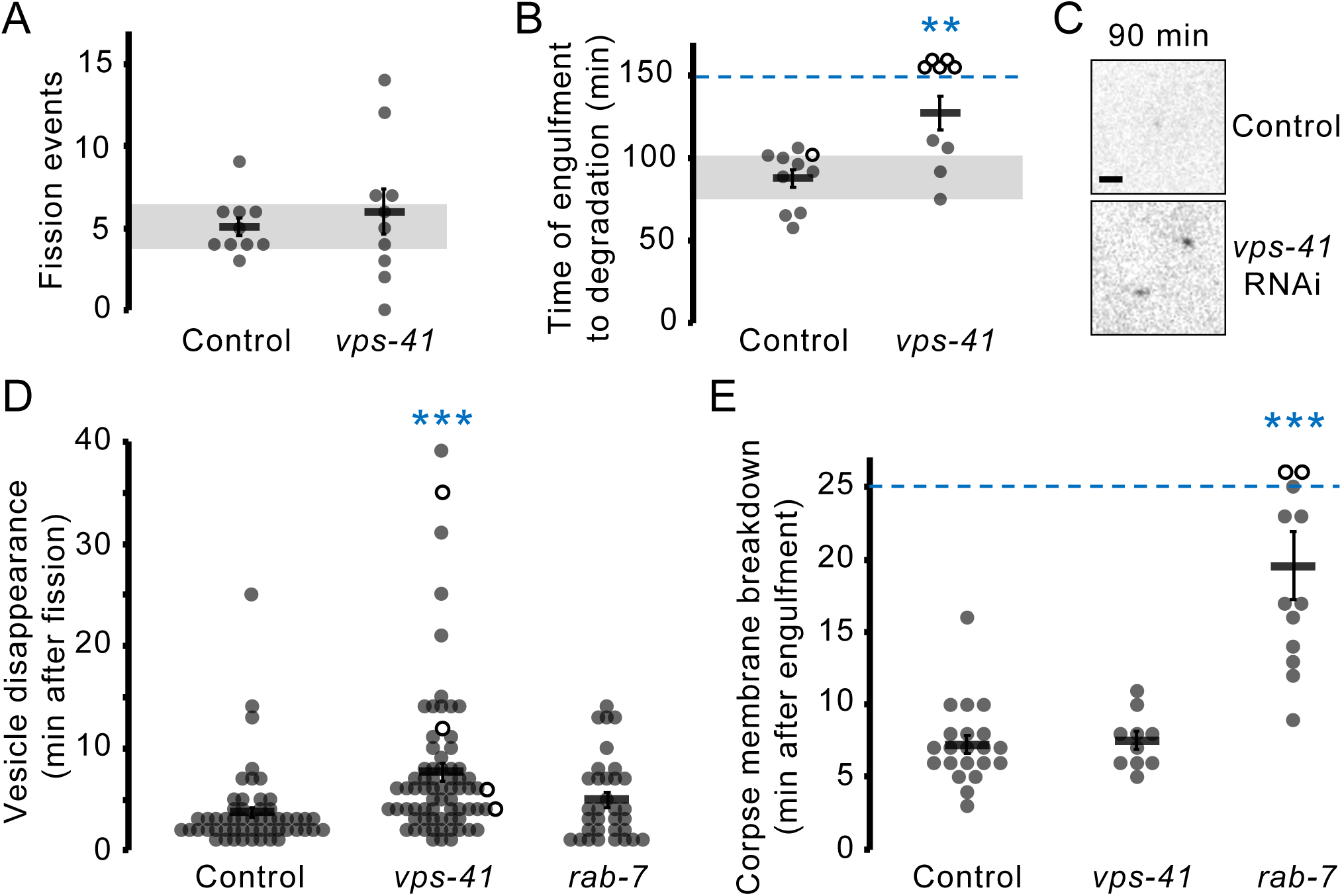
HOPS is required for degradation of small phagolysosomal vesicles. A) The number of phagolysosomal fission events was not affected after *vps-41* RNAi (6±1) compared to control embryos (5±1). B) Treatment with *vps-41* RNAi significantly delayed phagolysosome cargo degradation (127±11 min) compared to controls (88±6 min). Open circles denote the last frame of a time-lapse series in which the polar body phagolysosome did not disappear. Mean ± SEM is shown. Disappearance times beyond 150 min after engulfment are grouped above the dashed blue line. Grey bars show standard deviation of the mean in controls. **p<0.01 compared to control embryos using Student’s t-test. C) Inverted images of corpse phagolysosomes 90 minutes after engulfment in control and *vps-41* knockdown embryos. Scale bar is 2 μm. D) Phagolysosomal vesicles disappear quickly in control embryos (4±1 min) after fission from the phagolysosome, but last twice as long after knockdown of the HOPS subunit *vps-41* (8±1 min). Small vesicles disappear with normal timing after *rab-7* RNAi (5±1 min). Open circles denote the last frame of a time-lapse series in which the vesicle did not disappear. ***p<0.001 compared to control embryos using Student’s t-test with Bonferroni correction. E) Breakdown of the polar body corpse membrane inside the nascent phagolysosome takes 7±1 min in controls. Corpse membrane breakdown is delayed after *rab-* 7 knockdown (20±2 min) (Fazeli et al. 2018), but not affected after *vps-*41 knockdown (8±1 min). ***p<0.001 compared to control embryos using Student’s t-test with Bonferroni correction.

## Discussion

We used an *in vivo* system to observe and analyze the dynamics of single phagolysosomes containing a physiological cargo and uncovered the timing and molecular mechanisms of phagolysosome resolution. We previously showed that digestion of corpse cargo by lysosomal hydrolases starts after fusion of phagosomes to lysosomes and after breakdown of the corpse membrane inside the phagolysosome (Fazeli et al. 2018). Degradation of cargo proteins by lysosomal hydrolases generates free amino acids within the phagolysosome that are then exported by solute carriers, including SLC-36.1, to recruit and activate TORC1 on the phagolysosome surface (Fig. 9A-B). Independent of TORC1, BORC recruits the small GTPase ARL-8 to the phagolysosome (Fig. 9B), likely in its nucleotide-free form (Niwa et al. 2017). Based on the ability of SAM-4 to promote GTP exchange (Niwa et al. 2017), we predict that downstream of TORC1, BORC promotes GTP binding of ARL-8 (Fig. 9B). ARL-8 then needs to cycle between a GDP-, GTP-, and unbound state to dynamically link the phagolysosomal membrane to kinesin-1 to extend tubules and release phagolysosomal vesicles (Fig. 9C). These vesicles then depend on lysosomal fusion mediated by the HOPS tethering complex to facilitate cargo degradation and compartment resolution (Fig. 9D).

**Fig. 9:**
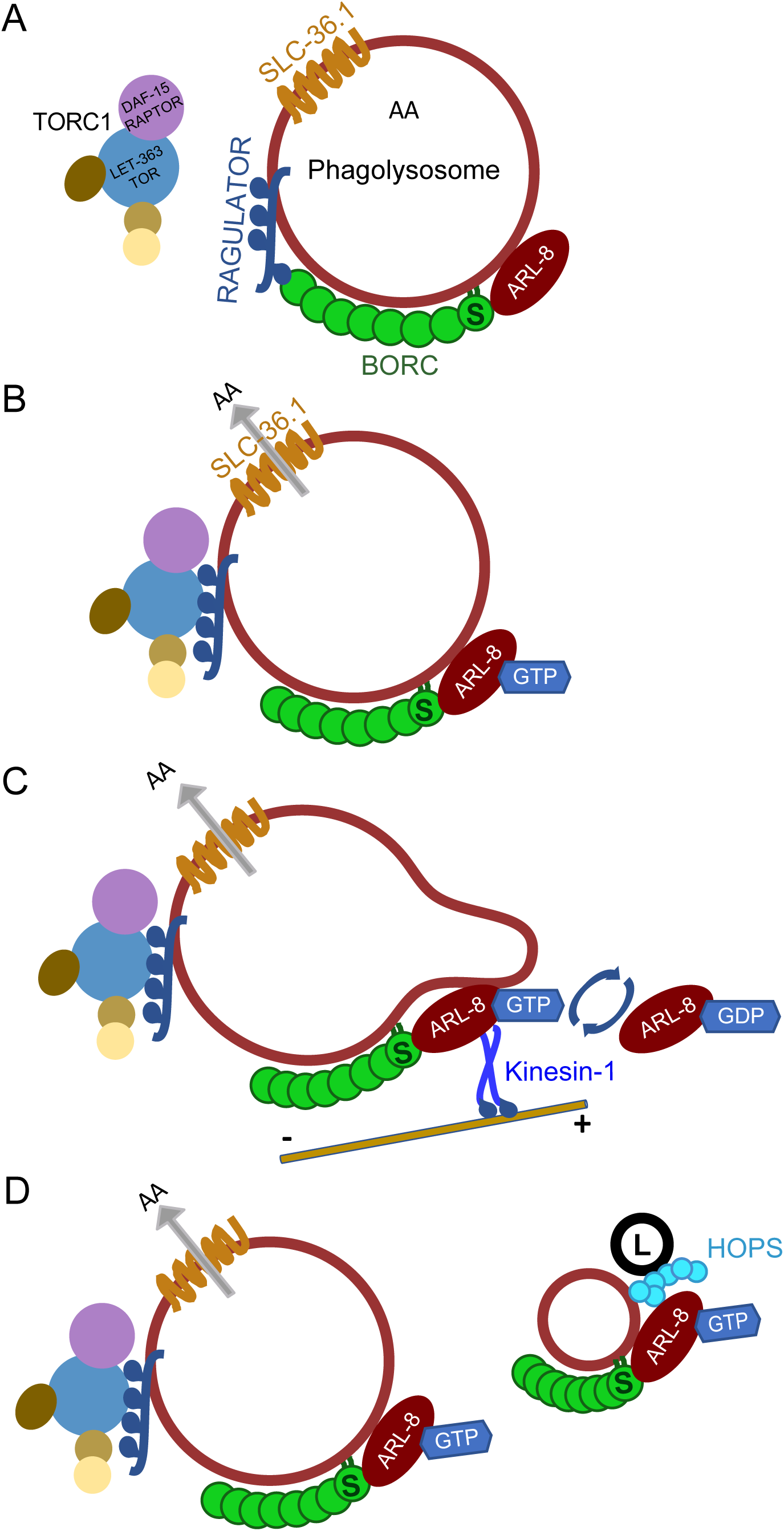
Model of phagolysosome vesiculation. A) After phagosome-lysosome fusion, phagolysosomes are decorated with lysosomal membrane proteins, including solute carriers, Ragulator, and BORC. The BORC subunit SAM- 4 (S) recruits nucleotide-free ARL-8 to the phagolysosome membrane. Digestion of corpse cargo by lysosomal hydrolases leads to amino acids in the phagolysosome lumen. B) Amino acids are exported by solute carriers, including SLC-36.1, and activate TORC1 to release BORC from Ragulator. SAM-4 promotes GTP-binding by ARL-8. C) ARL-8 needs to cycle between a GDP- and GTP-bound state to maintain its localization and promote tubulation via kinesin-1. D) Small phagolysosomal vesicles then fuse with lysosomes (L) using the HOPS tethering complex to facilitate rapid cargo degradation and resolution.

We discovered that similar TORC1-BORC-ARL8 machineries are involved in phagolysosome tubulation to lysosome trafficking (Table S1) (Pu et al. 2015; Farias et al. 2017; Niwa et al. 2017; De Pace et al. 2020). One possibility to explain the distinction between lysosomal movement and tubulation are that forces pull the phagolysosome in opposite directions, which could be accomplished by different microtubule motors. Indeed, phagosomes were recently shown to interact with minus-end-directed dynein motors via ARL8 (Keren-Kaplan et al. 2022), in addition to plus-end-directed kinesins. Alternatively, an organelle could hold the phagolysosome to prevent its movement and allow microtubule motors to instead drive tubulation. One intriguing possibility is that the newly discovered ER-phagolysosome contacts hold the phagolysosome in place to enable tubulation (Levin-Konigsberg et al. 2019). Although we show that BORC and ARL-8 act downstream of TORC1 in phagolysosomal tubulation, we did not observe mislocalization of ARL-8 in embryos deficient for amino acid transporters, Ragulator, or TORC1. In the absence of nutrients, LAMTOR/Ragulator is thought to keep BORC and ARL8 away from kinesin motor proteins that transport lysosomes towards the cell surface (Filipek et al. 2017; Pu et al. 2017), but not to influence ARL8 localization to the lysosome. Our data show that the mere release of BORC from Ragulator in *lmtr-2* mutants is not sufficient to allow ARL-8-mediated phagolysosome tubulation, contradicting the idea that the sole role of mTORC1 activation in tubulation is to release BORC inhibition. As LAMTOR/Ragulator also acts as a GEF for Rag GTPases, which in turn recruit mTORC1 to lysosomes (Sancak et al. 2010; Bar-Peled et al. 2012), *lmtr-2* mutants are predicted to not only fail to inhibit BORC, but also fail to recruit TORC1 to lysosomes, supporting the idea that TORC1 activation has an as yet unidentified role in phagolysosome tubulation beyond relieving BORC inhibition or influencing ARL-8 localization. One possibility is that TORC1 can affect factors downstream of ARL-8 for phagolysosome vesiculation.

Small GTPase localization is often guided by a GEF localized to a specific membrane, which in turn recruits and promotes membrane association of the GTPase (Blumer et al. 2013), however we revealed that a GEF is not sufficient to localize ARL-8 to lysosomes. We observed that GTP-locked ARL-8(Q75L) localizes non-specifically to membrane compartments but fails to localize to lysosomes. This is likely due to SAM-4 binding exclusively to nucleotide-free ARL- 8 (Niwa et al. 2017) and being unable to recruit GTP-locked ARL-8(Q75L). Similarly, SAM-4 is unable to stably recruit ARL-8 G4 mutants to lysosomes, likely because nucleotide-binding of G4 mutants is weakened (McCray et al. 2010), and they cannot stably bind GTP or GDP. Thus, SAM-4 binding to nucleotide-free ARL-8 and the ability to remain GTP-bound may be necessary to stably and specifically localize ARL-8 to lysosomes and phagolysosomes.

Similarly, the early endosomal localization of ARL-8 suggests the presence of an early endosomal Arl GEF that promotes endosomal membrane binding. This GEF is unlikely to be BORC because we observed ARL-8 accumulating on enlarged RAB-5-positive endosomes in BORC mutants. RAB-5 endosomes are enlarged upon interfering with ARL-8-lysosome association or the GDP-GTP cycle of ARL-8, suggesting that ARL-8 may need to be turned over on early endosomes to allow endosome maturation. These giant early endosomes resemble those formed in GTP-locked Rab5(Q79L) mutants (Stenmark et al. 1994), which also stably localize to early endosomes and disrupt endosome maturation. Intriguingly, Sasaki *et al*. observed that cell corpse phagosomes colocalized with RAB-5 for slightly longer in strong *arl-8* loss-of-function mutants (Sasaki et al. 2013), suggesting that ARL-8 may also promote Rab turnover for the maturation of early phagosomes and early endosomes.

By studying single phagosomes over their lifetimes, we uncovered the conserved mechanisms of how large phagolysosomes tubulate to form smaller vesicles and facilitate degradation. The ARL-8 effector HOPS likely promotes degradation of phagolysosomal vesicles via its role in lysosome fusion (Nguyen and Yates 2021), suggesting that multiple rounds of lysosome fusion are required for degradation of large phagolysosomal cargo. This begins with RAB-7-mediated fusion of the large phagosome with lysosomes to allow the cargo membrane to be broken down (Fazeli et al. 2018) and continues with HOPS-mediated lysosome fusion of small phagolysosomes. It will be interesting to determine why different proteins are required for lysosome fusion to different vesicles containing similar cargo.

Given the difficulty in unambiguously tracking a single phagosome in mammalian cells, let alone mammals, clearance of the *C. elegans* polar body is a powerful system to better understand proteins involved in phagolysosome resolution. Previously, mTOR was shown to be required for fragmentation of phagocytic and entotic cargo in both *C. elegans* and mammals (Krajcovic et al. 2013; Fazeli et al. 2018). Here, we observed for the first time that disrupting BORC subunits prevented phagolysosome fragmentation in both worms and mammals. Thus, evolutionarily conserved mechanisms conduct phagolysosome resolution. Given the importance of effectively resolving phagolysosomes containing pathogens or cell corpses, we predict that this pathway has a critical impact on modulation of immune responses as well as metabolism.

## Methods

### Worm strains and maintenance

*Caenorhabditis elegans* strains were maintained at room temperature according to standard protocol (Brenner 1974). For a list of strains used in this study, see Table S2. ARL-8::mCitrine knock-in did not alter developmental timing, viability, or fertility. While ARL-8(T34N) mutants were also viable and fertile, the combination of the mCitrine tag with the T34N mutation led to maternal-effect embryonic lethality in ARL-8(T34N)::mCit worms, similar to the strong loss-of-function deletion allele *tm2504*, suggesting that both perturbations partially disrupt ARL-8 function. Worms were PCR genotyped using primers in Table S3. Restriction fragment length polymorphisms (RFLP) using the following restriction enzymes were performed: *arl-8(jpn1)* with MboII, *arl-8(syb3658D133A)* with NlaIV, *arl-8(wy270)* with EcoRI, *arl-8(syb2255Q75L)* with Hpy188I, *arl-8(syb2260T34N)* with BstXI, *blos-8(js1354)* with HinDIII, *blos-9(js1352)* with BclI, *cup-14(cd32)* with PstI, *kxd-1(js1356)* with Hpy188III. *glo-2* mutants were phenotyped on a DM5500 wide-field fluorescence microscope (Leica) for the absence of auto-fluorescent and birefringent gut granules. *unc-104(e1265)* mutants were phenotyped for severe paralysis.

### RNAi experiments

RNAi was performed by feeding dsRNA-expressing bacteria at 25°C from the L1 larval stage through adulthood (64-72 hours) according to established protocols (Fraser et al. 2000). RNAi constructs were published previously (*daf-15* and *rict-1* RNAi (Qi et al. 2017)) or were obtained from available libraries (Source BioScience, *let-363 (sjj2_B0261.2a), rab-7 (mv_E03C9.3)*, *rub-1 (sjj_Y56A3A.16), slc-36.1 (sjj_Y43F4B.7), unc-116 (sjj_R05D3.7), and vps-41 (sjj2_F32A6.3)*).

### Light Microscopy

Embryos dissected on a cover slip in M9 buffer were mounted on a slide on top of an agarose pad. Sixteen 1.2 µm step Z-stacks were acquired sequentially for (GFP and) mCherry and DIC every minute at room temperature using a DM5500 wide-field fluorescence microscope with a HC PL APO 40X 1.3 NA oil objective lens supplemented with a DFC365 FX CCD camera controlled by LAS AF software (Leica). Confocal images were obtained using a Leica SP5 confocal with a HCX PL APO CS 40.0x1.25-NA oil objective with PMT detectors. To image ARL-8::mCit localization, dissected embryos were transferred into egg salts in a 4- or 8- chamber glass bottom slide (ibidi), illuminated using a Tilt light sheet (Mizar), and imaged using an Axio Observer 7 (Zeiss) with a 40X 1.4 NA oil objective lens and ORCA-Fusion sCMOS camera (Hamamatsu) controlled by SlideBook software (3i).

### Image Analysis

Time-lapse series were analyzed using Imaris (Bitplane). Internalization was defined as the first frame where the polar body moves away from the plasma membrane, which is likely to closely reflect closure of the phagocytic cup. Vesiculation events were scored when the vesicles were clearly distinct from each other, likely underestimating budding events. Each vesicle was followed individually until disappearance. Degradation was defined as the last frame where mCh reporters were observed after examining the following five frames for mCh puncta reappearance. A maximum of 170 min past engulfment was considered as the end of each time lapse if the phagolysosome was not yet degraded.

### Colocalization analysis

z-stacks of 1-4-cell embryos where deconvolved and smoothened using a median filter to reduce the noise and the background was masked in Imaris. For each channel, surfaces were created to annotate the fluorescent structures using 3D View in Imaris. Thresholded colocalization of ARL-8::mCit with SAM-4::mSc or mCh::RAB-5 was measured using the Coloc tool of Imaris and percent of ARL-8 colocalization and Pearson coefficient in colocalized volume was reported.

### Antibody staining

Gravid worms were dissected in water on a coverslip to release embryos and transferred to 0.1% polylysine-coated slides and frozen on dry ice. Eggshells were cracked by flicking off the coverslip and embryos were fixed in methanol before staining. The following antibodies were used: mouse α-LMP-1 antibody (1:50, AB_2161795, Developmental Studies Hybridoma Bank), chicken α-GFP (1:500, AB_2307313, Aves). Slides were counterstained with DAPI to label DNA and mounted using DABCO.

### Image processing

For clarity, images were rotated, colorized and the intensity was adjusted using Adobe Photoshop. All images show a single optical section (Z), except Fig. 1D, where the last two panels are maximum projection of two Zs with 1.2 μm steps. Several Zs with 1.2 μm steps were maximum projected in Imaris for videos.

### Mammalian cell culture

J774 murine macrophages were cultured in DMEM supplemented with 100 U/ml penicillin, 100 µg/ml streptomycin, and 10% heat-inactivated fetal bovine serum. Cells were passaged by scraping when they reached 80 – 90% confluency.

### Generation of mutant macrophage clones

Cas9-expressing J774 cells were transduced with lentivirus based on pMCB320 (Addgene plasmid #89359) with mCherry replaced by BFP and containing the sgRNA sequences given in the table below. Two days after transduction, sgRNA-expressing cells were selected with puromycin (2 µg/ml). CRISPR-modified J774 cells were isolated to identify clonal lines.

Genomic DNA from clones was PCR amplified, sequenced, and subjected to TIDE analysis(Brinkman et al. 2014) to verify genome edits (Fig. S4).

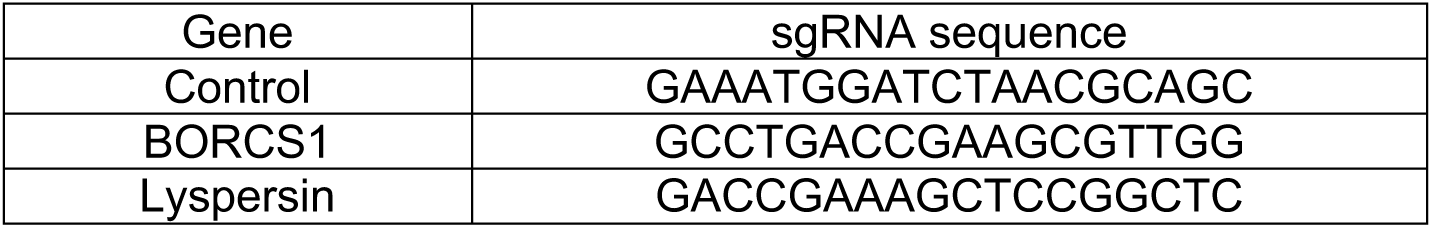

### Fragmentation assay

J774 cells were plated on coverslips in 12-well plates (75,000 cells/well) for 24 hours. Sheep red blood cells (SRBC) were opsonized with rabbit anti-sheep IgG (MP Biomedical) as described (Montano et al. 2018) and labeled with TAMRA-succinimidyl ester. Macrophages were fed IgG-coated SRBCs at a ratio of 3 SRBC per macrophage for 30 minutes, washed with D-PBS, placed in fresh culture media, and incubated at 37°C for 6 h. J774 cells were then washed with D-PBS and fixed with 2% PFA for 12 min before imaging on a confocal microscope. The extent of phagolysosome fragmentation was scored (Stage I-III) for >120 cells in three separate experiments using the same clonal cell lines.

### Statistical evaluation

Student’s one-tailed *t*-test, or Chi-squared test were used to test statistical significance using the Bonferroni correction to adjust for multiple comparisons. Mean ± standard error of the mean is depicted in graphs and text, except for Fig. 6D, where mean ± standard deviation is shown.

## Supplemental movie legends

Supplemental Movie 1. The corpse phagolysosome tubulates to release cargo- containing vesicles in a *C. elegans* embryo. Related to Figure 1.

The second polar body (arrowhead) is phagocytosed and the phagolysosome containing the corpse of the 2^nd^ polar body tubulates to release vesicles, most notably starting after 50 minutes. The phagolysosomal vesicles are then resolved by 1.5 hours. The embryo expresses the PI4,5P2 reporter mCh::PH::ZF1, which initially marks the plasma membrane in all cells, including the polar bodies, but is progressively degraded from somatic cells, starting with the anterior cells that phagocytose the second polar body. By 30 minutes, the mCh::PH::ZF1 cargo in the phagolysosome is visible without background labeling from the engulfing cell membrane. The inverted right panel shows the magnified region containing the polar body phagolysosome marked by the dashed rectangle. Time-lapse data were collected every minute and displayed at 6 fps. 10 Zs were projected (Z interval 1.2 μm).

**Supplemental Movie 2: The corpse phagolysosome fails to tubulate or release cargo-containing vesicles in a BORC mutant embryo.** Related to Figure 2.

The second polar body (arrowhead) is phagocytosed in a *blos-7/Lyspersin* mutant embryo, but the phagolysosome fails to tubulate or release vesicles, remaining large and bright even after 2.5 hours. The embryo expresses the PI4,5P2 reporter mCh::PH::ZF1, which initially marks the plasma membrane in all cells, including the polar bodies, but is progressively degraded from somatic cells, starting with the anterior cells that phagocytose the second polar body. See Supplemental Movie 1 for the control embryo. The inverted right panel shows the magnified region containing the polar body phagolysosome marked by the dashed rectangle. Time-lapse data were collected every minute and displayed at 6 fps. 10 Zs were projected (Z interval 1.2 μm).

## Acknowledgements

Anna Henrich, Martin Boos, Kaitlyn Spees, Jasmine Garcia and Heather Chorzempa provided technical assistance. The authors thank the imaging facility of the Rudolf Virchow Center for providing support for imaging and data analysis. Strains, constructs, and reagents were generously provided by Ralf Baumeister, Hanna Fares, Greg Hermann, Shinsuke Niwa, Michael Nonet, Karen Oegema, Kang Shen, Japan National Bioresource Project (NBRP), and the Caenorhabditis Genetics Center (CGC), which is funded by NIH Office of Research Infrastructure Programs (P40 OD010440). Todd Blankenship, Julia Frondoni, Shruti Kolli, and Alex Nguyen provided valuable comments on the manuscript. This work was funded by Deutsche Forschungsgemeinschaft (DFG) grants FA1046/3-1 to GF and STI700/1-1 to CS.

## Author Contributions

GF designed, performed, analyzed, and supervised most worm experiments and wrote the manuscript together with RLK and AMW. RLK designed, performed, and analyzed mammalian CRISPR experiments. MCB and CS supervised experiments. AMW designed and supervised experiments, performed light sheet microscopy, and analyzed data.

---

**Supplemental Fig. 1:**
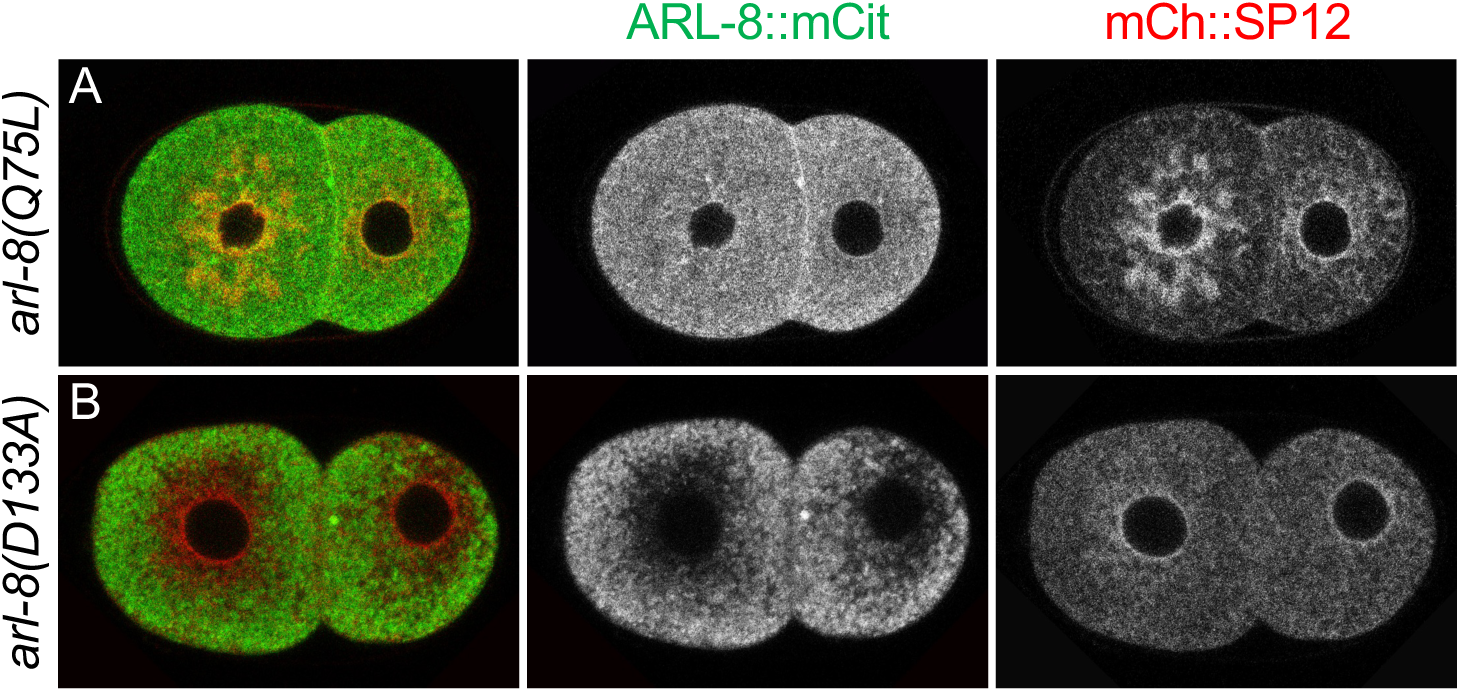
GTP-locked ARL-8, but not unlocked ARL-8, localizes to ER. A) The GTP-locked *arl-8(Q75L)*::mCit allele (green) localizes to membrane compartments including plasma membrane and ER marked by mCh::SP12 (red). B) The fast exchange G4 mutant *arl-8(D133A)*::mCit allele (green) is excluded from the plasma membrane and ER marked by mCh::SP12 (red).

**Supplemental Fig. 2:**
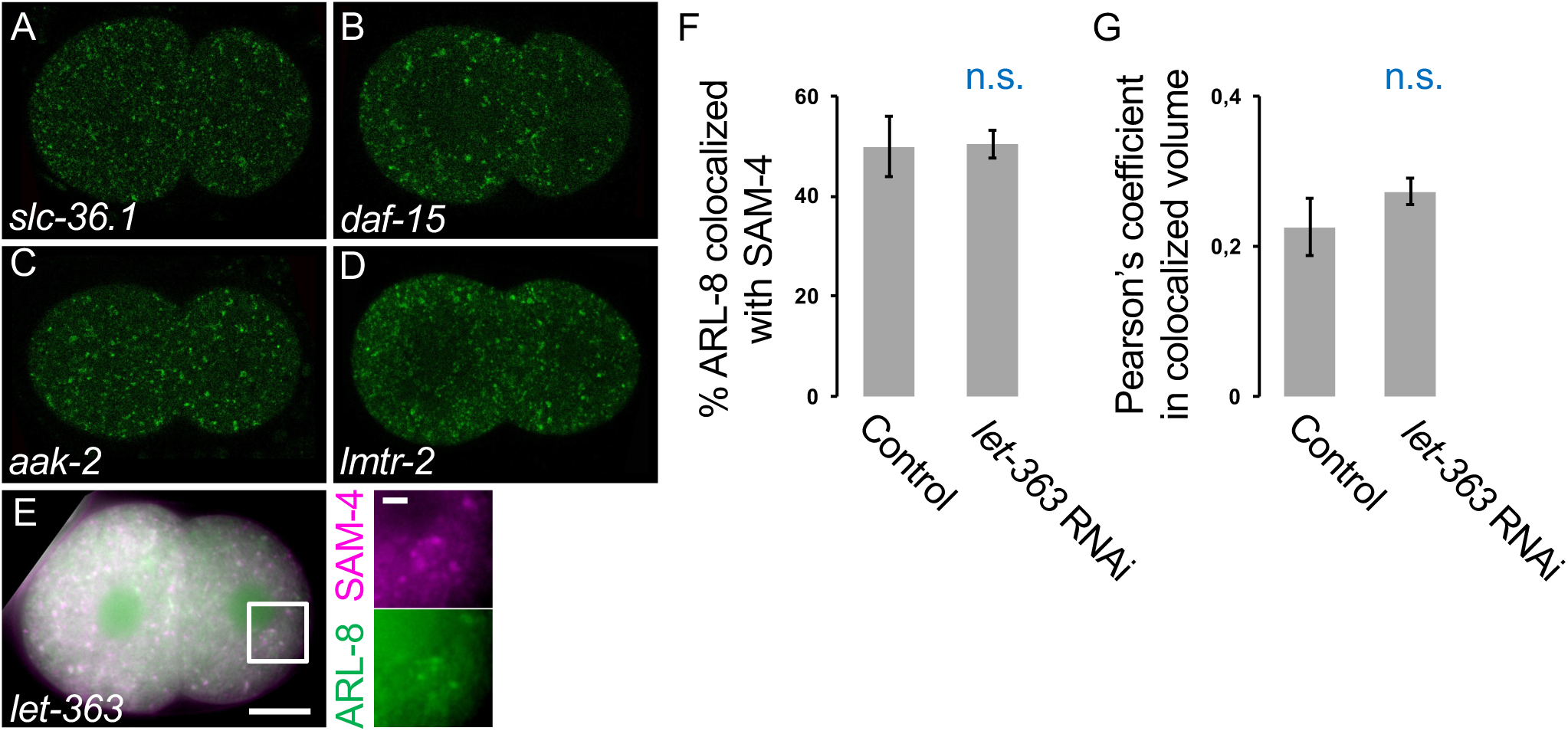
TORC1 is not required for ARL-8 localization. A-D) Endogenously-expressed mCit::ARL*-8* localizes to discrete puncta (lysosomes and endosomes) in embryos treated with RNAi against *slc-36.1* (A) or *Raptor/daf-15* (B), similar to control embryos (see Fig. 4B, G & K). Deletion mutations in AMPK subunit *aak-2(ok524)* (C) or Ragulator subunit *lmtr-2(tm2367)* (D) also did not change the localization of ARL-8. E) ARL- 8::mCit still colocalizes with the lysosomal resident BORC subunit SAM-4 after RNAi treatment against *TOR/let-363*. Scale bars are 10 μm in the main panel and 2 μm in insets. F-G) mCit::ARL-8 colocalization with lysosome resident SAM-4::mSc did not significantly differ after treatment with *let-363* RNAi judged by percentage of colocalization (F) or Pearson’s coefficient (G).

**Supplemental Fig. 3:**
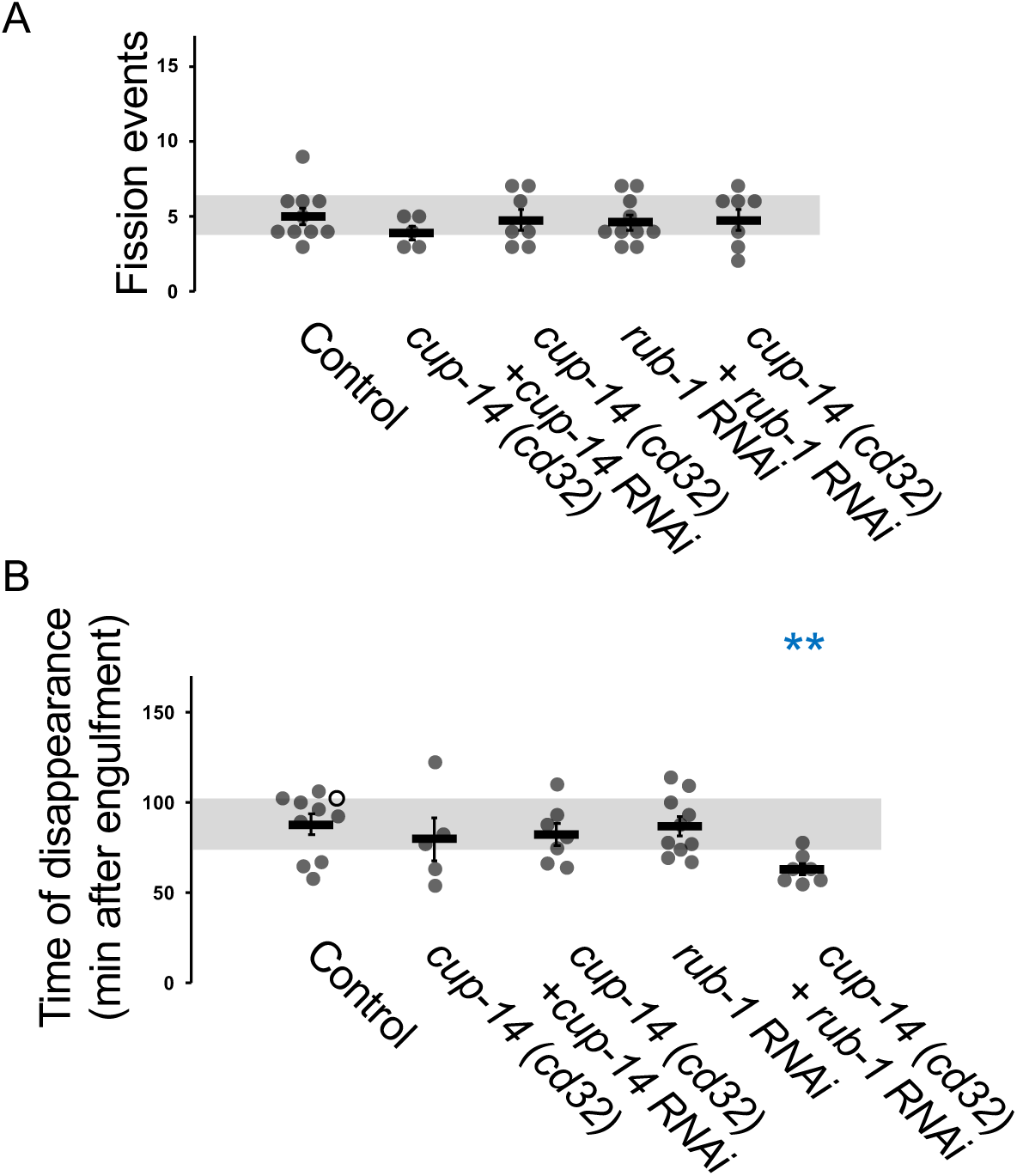
Disrupting PLEKHM proteins does not alter vesiculation or degradation. A) Phagolysosomal fission events were not affected compared to control embryos (5±1) after disrupting PLEKHM family proteins in *cup-14(cd*32) mutants with (5±1) or without (4±0) *cup- 14 RNAi* or after *rub-1* RNAi treatment in a control (5±0) or *cup-14(cd32)* mutant background (5±1). B) The 2^nd^ polar body cargo mCh::PH::ZF1 disappears from the phagolysosome 88±6 min after internalization. Disappearance of the phagolysosome cargo was not affected in *cup- 14(cd32)* mutants with (82±6) or without (80±12) *cup-14 RNAi* or after control worms were treated with *rub-1* RNAi (87±5). Treatment of *cup-14(cd32)* mutants with *rub-1* RNAi accelerated phagolysosome degradation (63±3 min). Open circles denote the last frame of a time-lapse series in which the polar body phagolysosome did not disappear. Mean ± SEM is shown. Grey bars show standard deviation of the mean in controls. **p<0.01 compared to control embryos using Student’s t-test with Bonferroni correction.

**Supplemental Fig. 4:**
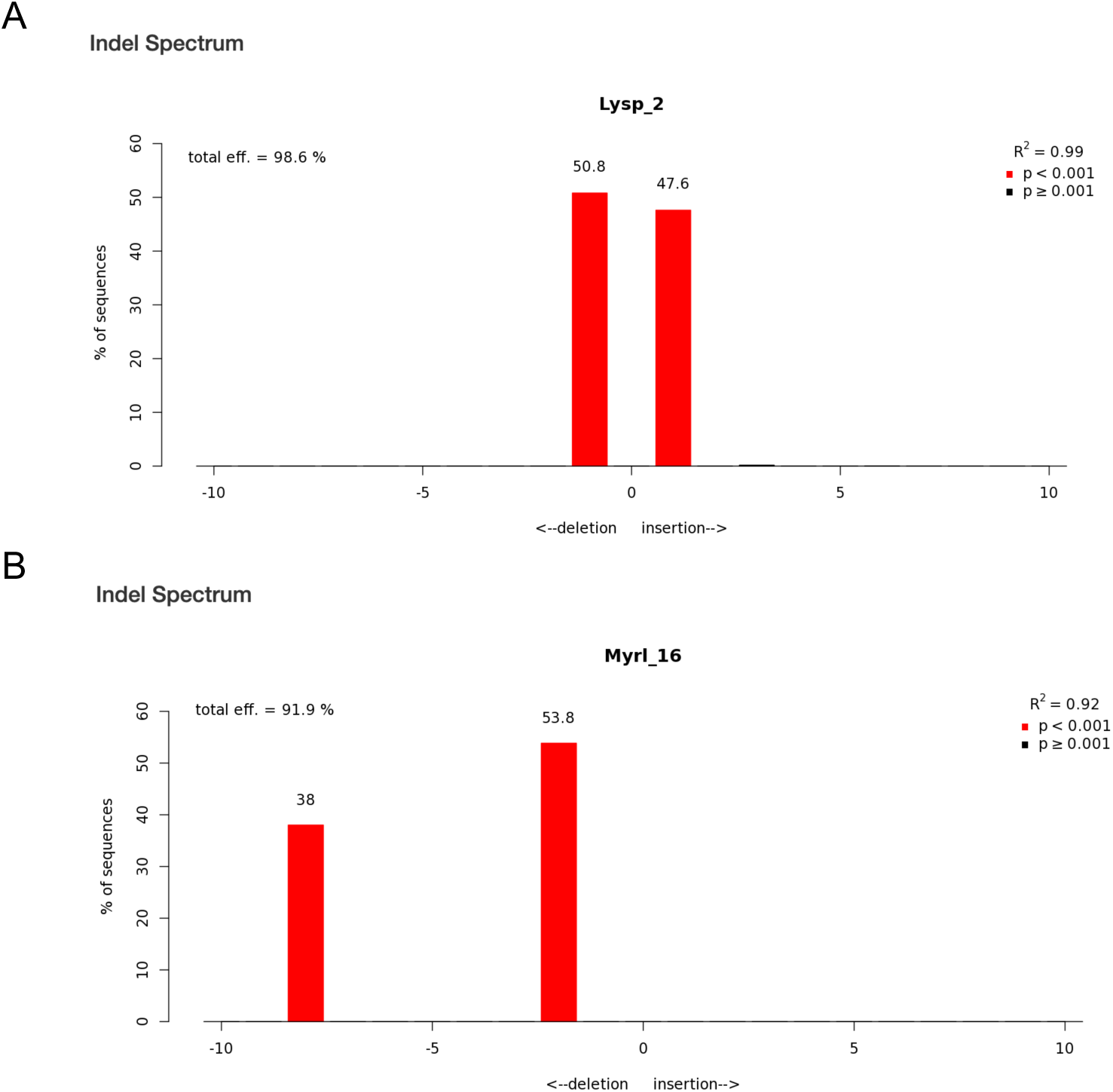
Verification of mutant macrophage cell lines. A-B) TIDE analysis of CRISPR/Cas9 editing of Lyspersin (A), or Myrlysin (B) mutant clones. The x axis indicates the number of bases inserted or deleted to induce frame shift mutations. Both cell lines are transheterozygous for two indels.

**Table S1:**
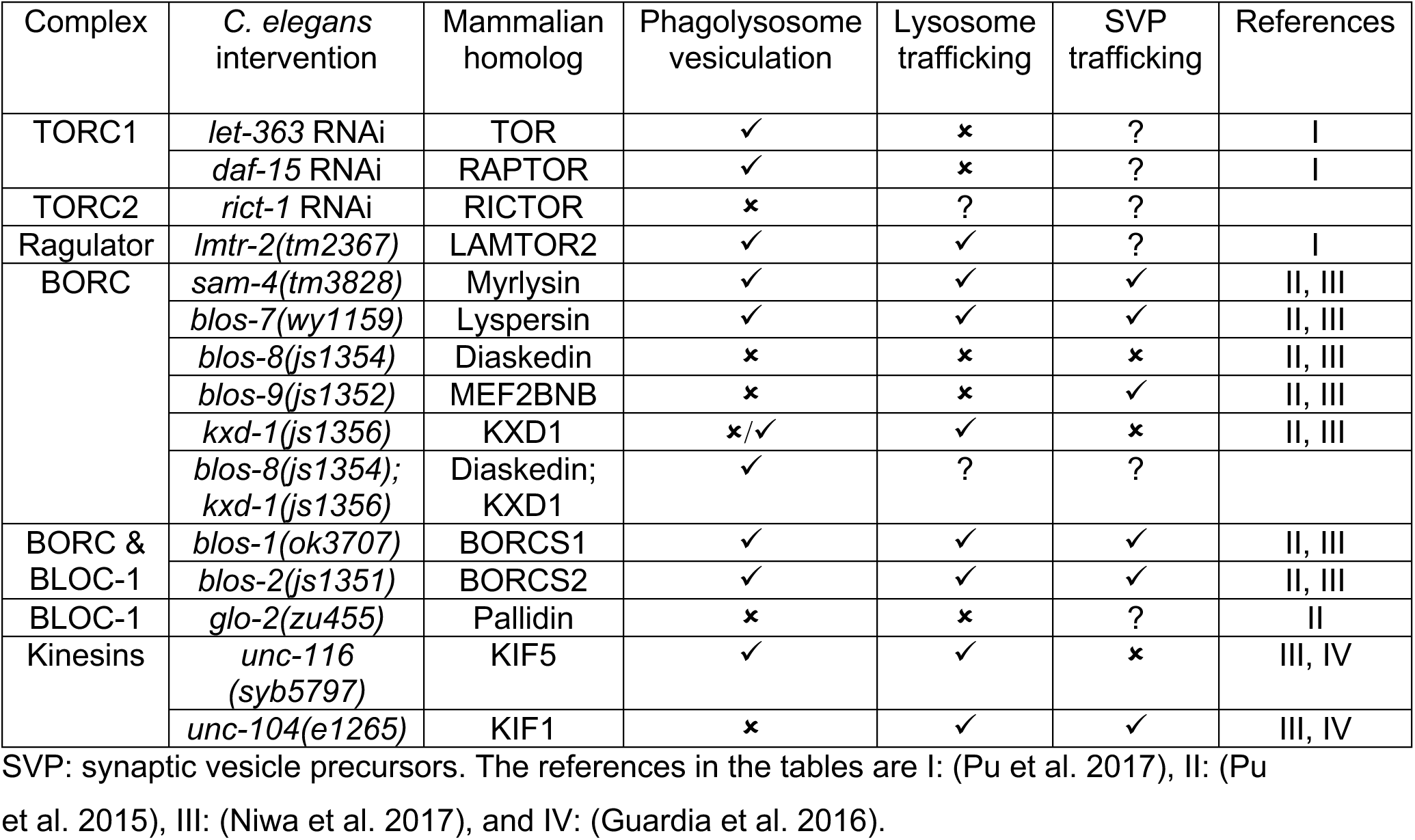
Summary of the known roles of protein complexes during phagolysosome vesiculation or organelle trafficking.

| Complex | *C. elegans* intervention | Mammalian homolog | Phagolysosome vesiculation | Lysosome trafficking | SVP trafficking | References |
| --- | --- | --- | --- | --- | --- | --- |
| TORC1 | *let-363* RNAi | TOR | ✓ | ✗ | ? | I |
|  | *daf-15* RNAi | RAPTOR | ✓ | ✗ | ? | I |
| TORC2 | *rict-1* RNAi | RICTOR | ✗ | ? | ? |  |
| Ragulator | *lmtr-2(tm2367)* | LAMTOR2 | ✓ | ✓ | ? | I |
| BORC | *sam-4(tm3828)* | Myrlysin | ✓ | ✓ | ✓ | II, III |
|  | *blos-7(wy1159)* | Lyspersin | ✓ | ✓ | ✓ | II, III |
|  | *blos-8(js1354)* | Diaskedin | ✗ | ✗ | ✗ | II, III |
|  | *blos-9(js1352)* | MEF2BNB | ✗ | ✗ | ✓ | II, III |
|  | *kxd-1(js1356)* | KXD1 | ✗/✓ | ✓ | ✗ | II, III |
|  | *blos-8(js1354); kxd-1(js1356)* | Diaskedin; KXD1 | ✓ | ? | ? |  |
| BORC & BLOC-1 | *blos-1(ok3707)* | BORCS1 | ✓ | ✓ | ✓ | II, III |
|  | *blos-2(js1351)* | BORCS2 | ✓ | ✓ | ✓ | II, III |
| BLOC-1 | *glo-2(zu455)* | Pallidin | ✗ | ✗ | ? | II |
| Kinesins | *unc-116 (syb5797)* | KIF5 | ✓ | ✓ | ✗ | III, IV |
|  | *unc-104(e1265)* | KIF1 | ✗ | ✓ | ✓ | III, IV |

SVP: synaptic vesicle precursors. The references in the tables are I: (Pu et al. 2017), II: (Pu et al. 2015), III: (Niwa et al. 2017), and IV: (Guardia et al. 2016).

**Table S2:**
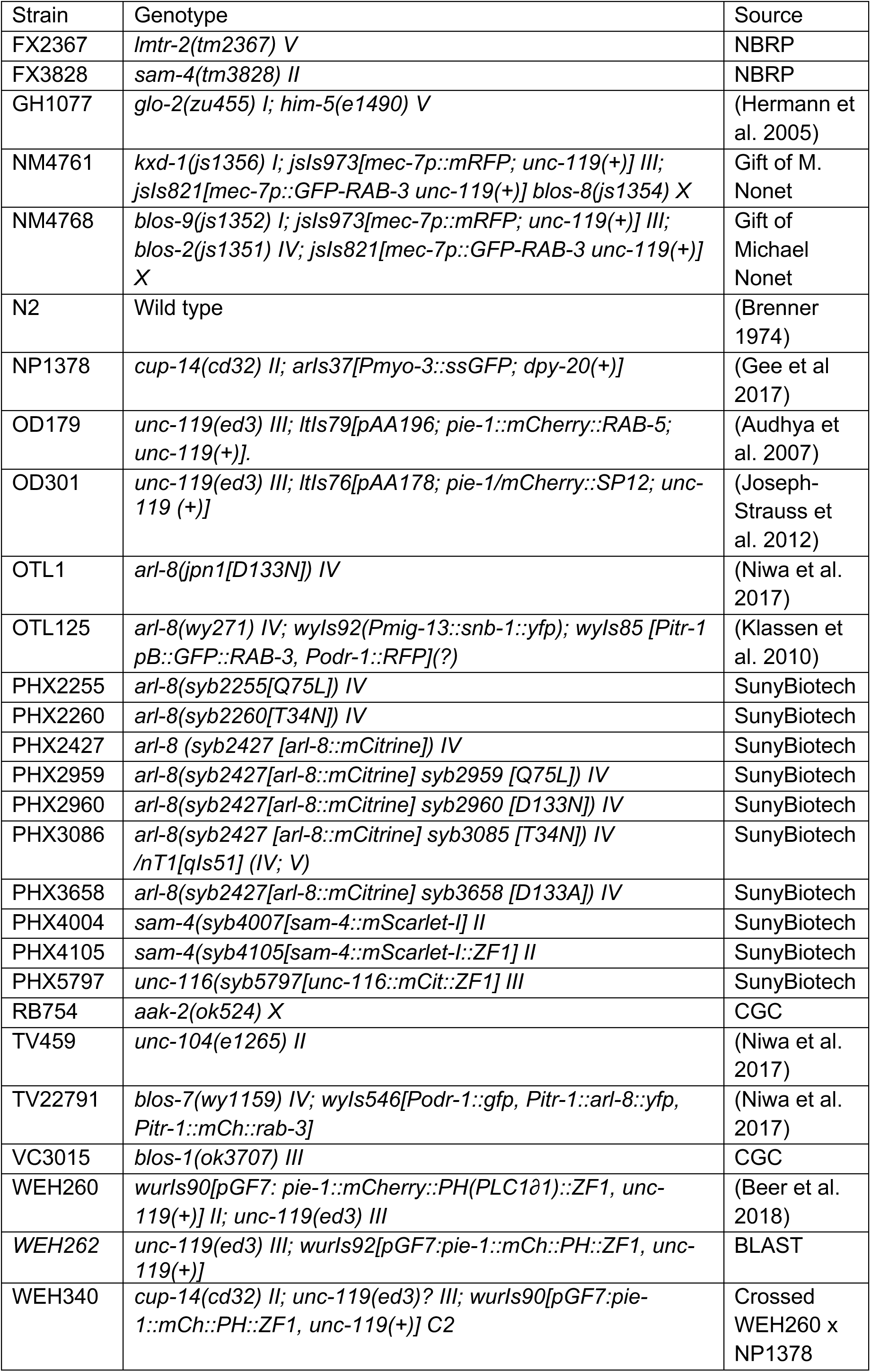
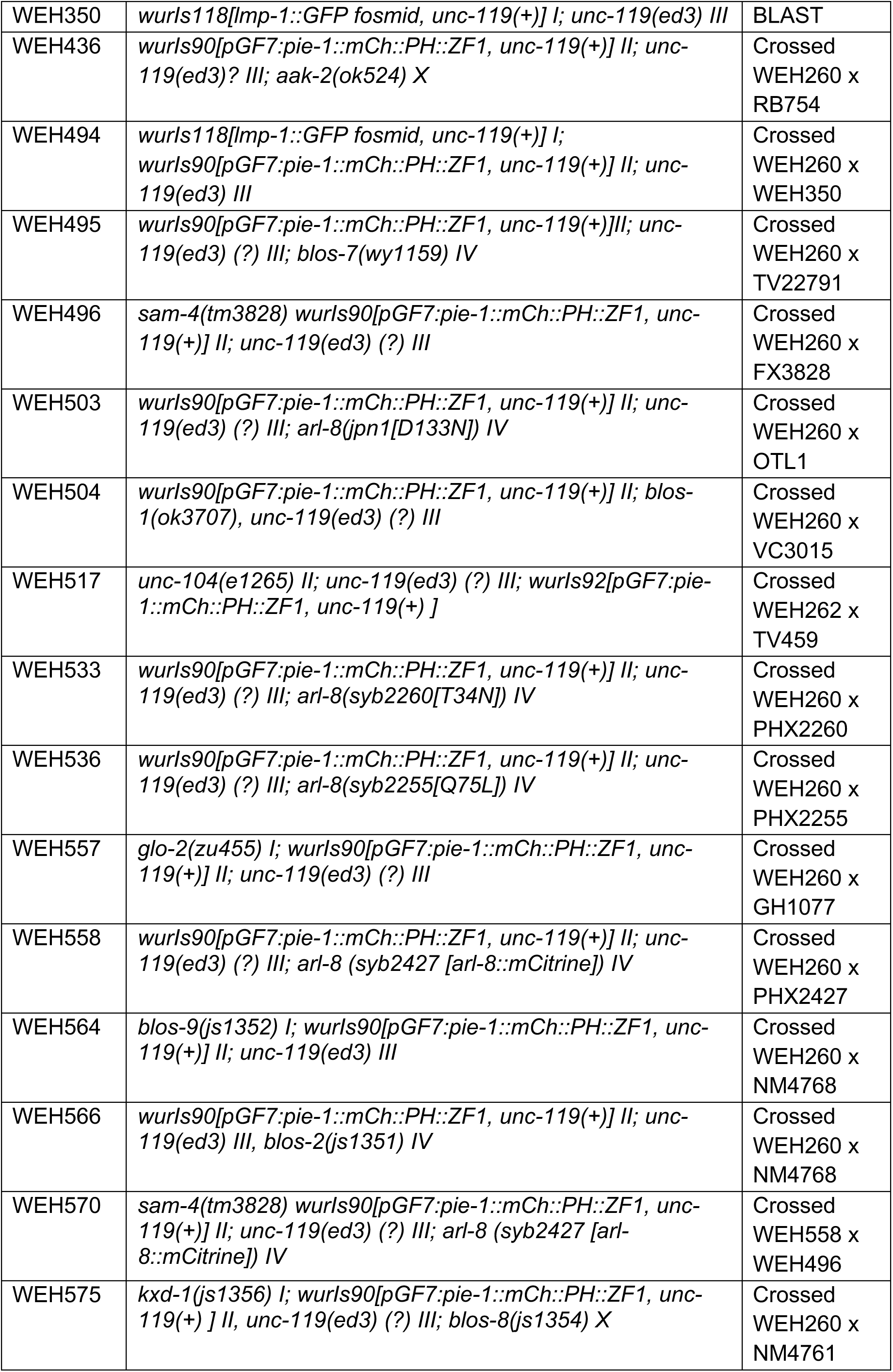
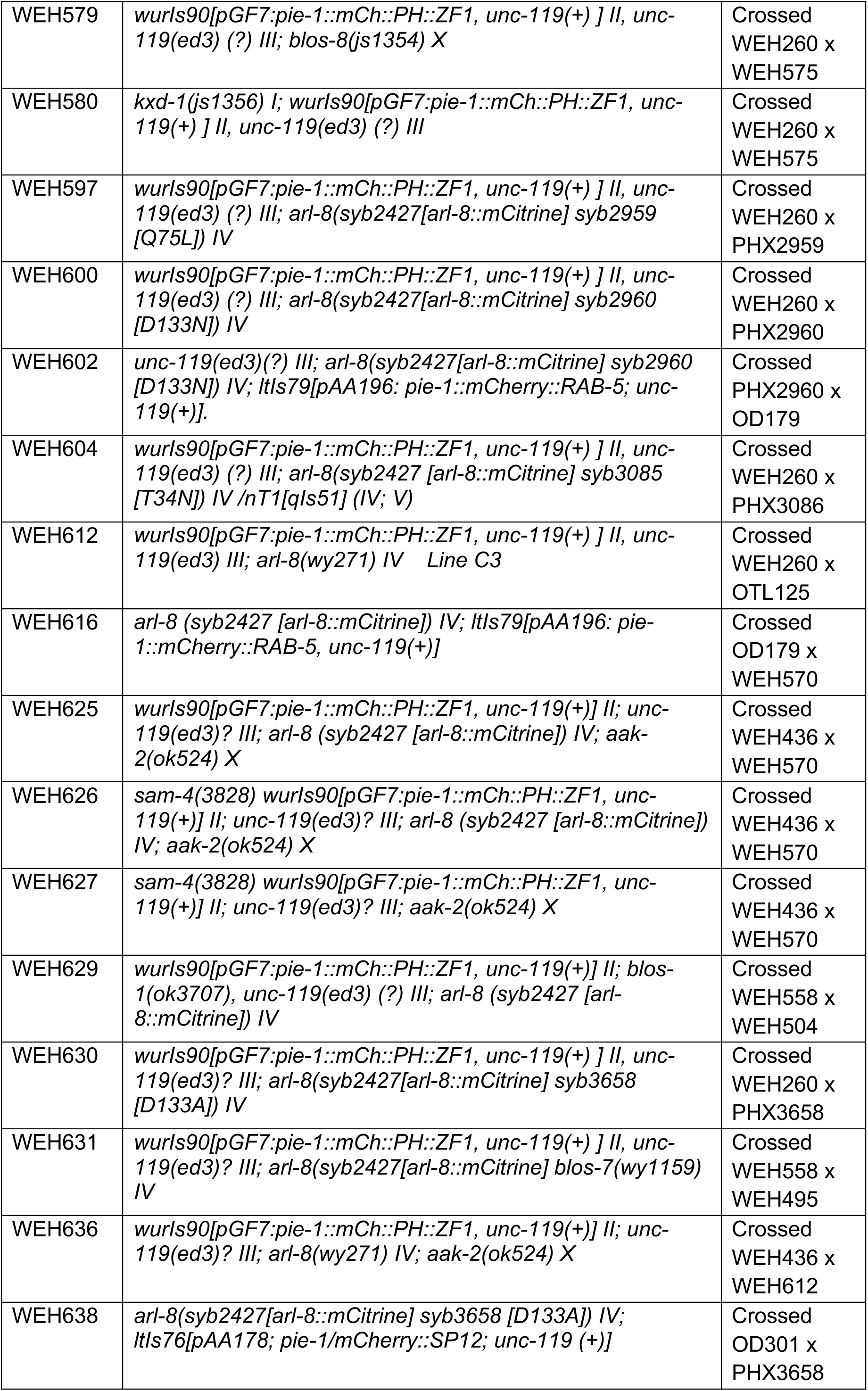
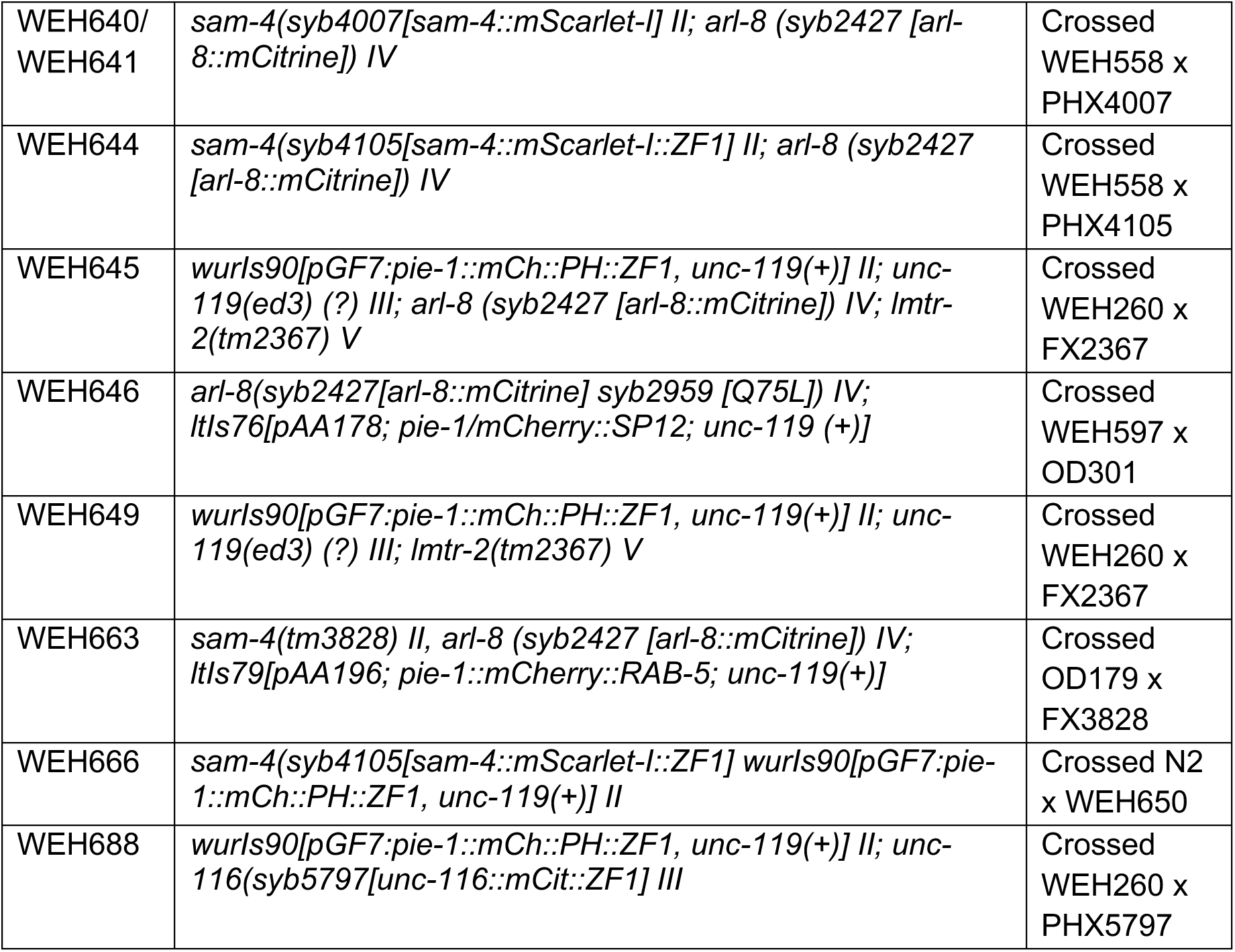
Strains used in this study.

**Table S3:**
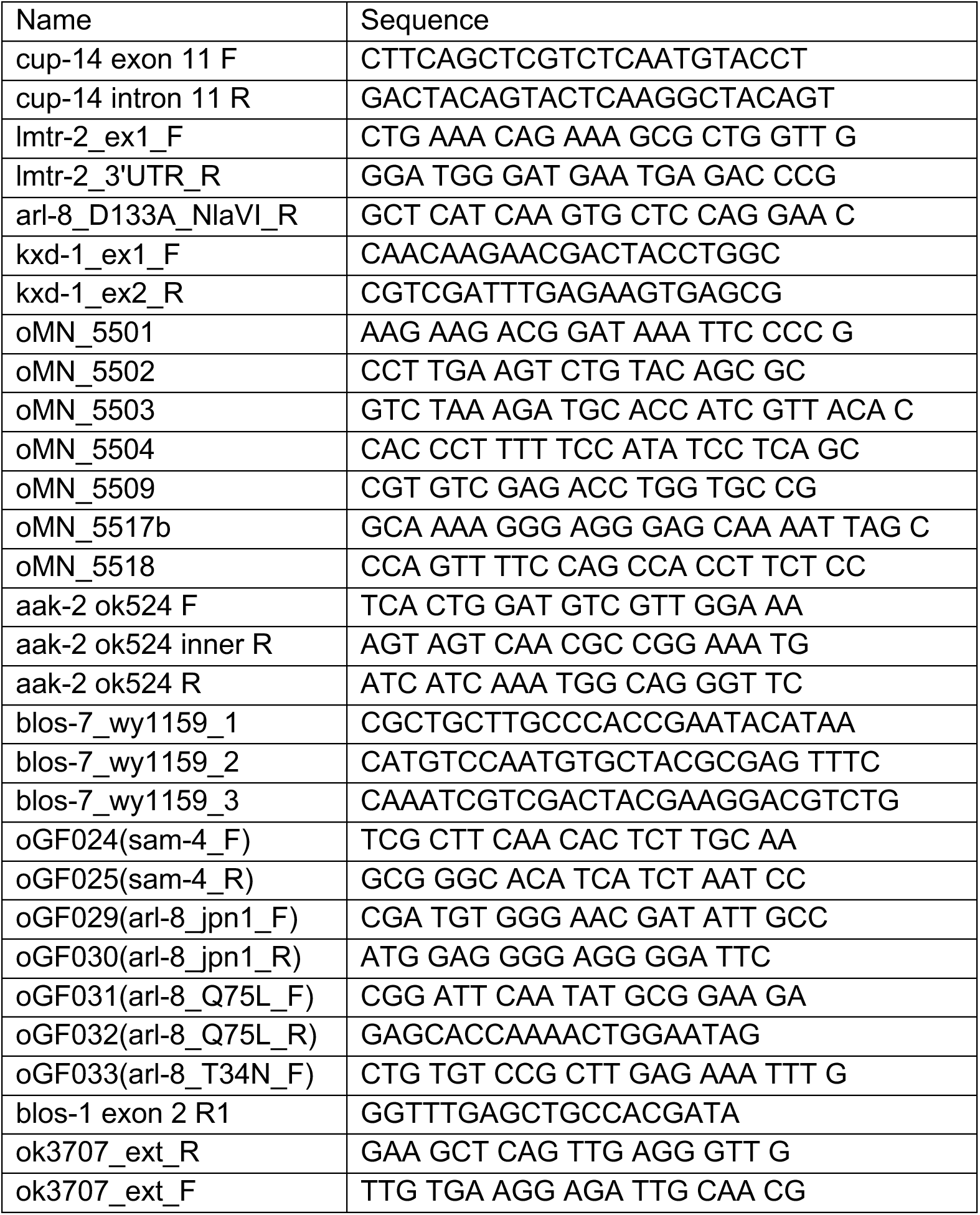
Primers used in this study.

## References

Armenti ST, Lohmer LL, Sherwood DR, Nance J. 2014. Repurposing an endogenous degradation system for rapid and targeted depletion of C. elegans proteins. Development 141: 4640–4647.

Audhya A, McLeod IX, Yates JR, Oegema K. 2007. MVB-12, a fourth subunit of metazoan ESCRT-I, functions in receptor downregulation. PLoS One 2: e956.

Bagshaw RD, Callahan JW, Mahuran DJ. 2006. The Arf-family protein, Arl8b, is involved in the spatial distribution of lysosomes. Biochem Biophys Res Commun 344: 1186–1191.

Bar-Peled L, Schweitzer LD, Zoncu R, Sabatini DM. 2012. Ragulator is a GEF for the rag GTPases that signal amino acid levels to mTORC1. Cell 150: 1196–1208.

Beer KB, Rivas-Castillo J, Kuhn K, Fazeli G, Karmann B, Nance JF, Stigloher C, Wehman AM. 2018. Extracellular vesicle budding is inhibited by redundant regulators of TAT-5 flippase localization and phospholipid asymmetry. Proc Natl Acad Sci U S A 115: E1127–E1136.

Blumer J, Rey J, Dehmelt L, Mazel T, Wu YW, Bastiaens P, Goody RS, Itzen A. 2013. RabGEFs are a major determinant for specific Rab membrane targeting. J Cell Biol 200: 287–300.

Brenner S. 1974. The genetics of Caenorhabditis elegans. Genetics 77: 71–94.

Brinkman EK, Chen T, Amendola M, van Steensel B. 2014. Easy quantitative assessment of genome editing by sequence trace decomposition. Nucleic Acids Res 42: e168.

Cherfils J, Zeghouf M. 2013. Regulation of small GTPases by GEFs, GAPs, and GDIs. Physiol Rev 93: 269–309.

Colaco A, Jaattela M. 2017. Ragulator-a multifaceted regulator of lysosomal signaling and trafficking. J Cell Biol 216: 3895–3898.

De Pace R, Britt DJ, Mercurio J, Foster AM, Djavaherian L, Hoffmann V, Abebe D, Bonifacino JS. 2020. Synaptic Vesicle Precursors and Lysosomes Are Transported by Different Mechanisms in the Axon of Mammalian Neurons. Cell Rep 31: 107775.

Farias GG, Guardia CM, De Pace R, Britt DJ, Bonifacino JS. 2017. BORC/kinesin-1 ensemble drives polarized transport of lysosomes into the axon. Proc Natl Acad Sci U S A 114: E2955–E2964.

Fazeli G, Stetter M, Lisack JN, Wehman AM. 2018. C. elegans Blastomeres Clear the Corpse of the Second Polar Body by LC3-Associated Phagocytosis. Cell Rep 23: 2070–2082.

Filipek PA, de Araujo MEG, Vogel GF, De Smet CH, Eberharter D, Rebsamen M, Rudashevskaya EL, Kremser L, Yordanov T, Tschaikner P et al. 2017. LAMTOR/Ragulator is a negative regulator of Arl8b- and BORC-dependent late endosomal positioning. J Cell Biol 216: 4199–4215.

Fond AM, Ravichandran KS. 2016. Clearance of Dying Cells by Phagocytes: Mechanisms and Implications for Disease Pathogenesis. Adv Exp Med Biol 930: 25–49.

Fraser AG, Kamath RS, Zipperlen P, Martinez-Campos M, Sohrmann M, Ahringer J. 2000. Functional genomic analysis of C. elegans chromosome I by systematic RNA interference. Nature 408: 325–330.

Gan Q, Wang X, Zhang Q, Yin Q, Jian Y, Liu Y, Xuan N, Li J, Zhou J, Liu K et al. 2019. The amino acid transporter SLC-36.1 cooperates with PtdIns3P 5-kinase to control phagocytic lysosome reformation. J Cell Biol 218: 2619–2637.

Ghose P, Wehman AM. 2021. The developmental and physiological roles of phagocytosis in Caenorhabditis elegans. Curr Top Dev Biol 144: 409–432.

Guardia CM, Farias GG, Jia R, Pu J, Bonifacino JS. 2016. BORC Functions Upstream of Kinesins 1 and 3 to Coordinate Regional Movement of Lysosomes along Different Microtubule Tracks. Cell Rep 17: 1950–1961.

Hermann GJ, Schroeder LK, Hieb CA, Kershner AM, Rabbitts BM, Fonarev P, Grant BD, Priess JR. 2005. Genetic analysis of lysosomal trafficking in Caenorhabditis elegans. Mol Biol Cell 16: 3273–3288.

Joseph-Strauss D, Gorjanacz M, Santarella-Mellwig R, Voronina E, Audhya A, Cohen-Fix O. 2012. Sm protein down-regulation leads to defects in nuclear pore complex disassembly and distribution in C. elegans embryos. Dev Biol 365: 445–457.

Keren-Kaplan T, Saric A, Ghosh S, Williamson CD, Jia R, Li Y, Bonifacino JS. 2022. RUFY3 and RUFY4 are ARL8 effectors that promote coupling of endolysosomes to dynein-dynactin. Nat Commun 13: 1506.

Khatter D, Raina VB, Dwivedi D, Sindhwani A, Bahl S, Sharma M. 2015a. The small GTPase Arl8b regulates assembly of the mammalian HOPS complex on lysosomes. J Cell Sci 128: 1746–1761.

Khatter D, Sindhwani A, Sharma M. 2015b. Arf-like GTPase Arl8: Moving from the periphery to the center of lysosomal biology. Cell Logist 5: e1086501.

Kim J, Kundu M, Viollet B, Guan KL. 2011. AMPK and mTOR regulate autophagy through direct phosphorylation of Ulk1. Nat Cell Biol 13: 132–141.

Klassen MP, Wu YE, Maeder CI, Nakae I, Cueva JG, Lehrman EK, Tada M, Gengyo-Ando K, Wang GJ, Goodman M et al. 2010. An Arf-like small G protein, ARL-8, promotes the axonal transport of presynaptic cargoes by suppressing vesicle aggregation. Neuron 66: 710–723.

Krajcovic M, Krishna S, Akkari L, Joyce JA, Overholtzer M. 2013. mTOR regulates phagosome and entotic vacuole fission. Mol Biol Cell 24: 3736–3745.

Lancaster CE, Fountain A, Dayam RM, Somerville E, Sheth J, Jacobelli V, Somerville A, Terebiznik MR, Botelho RJ. 2021. Phagosome resolution regenerates lysosomes and maintains the degradative capacity in phagocytes. J Cell Biol 220.

Langemeyer L, Ungermann C. 2015. BORC and BLOC-1: Shared subunits in trafficking complexes. Dev Cell 33: 121–122.

Levin-Konigsberg R, Montano-Rendon F, Keren-Kaplan T, Li R, Ego B, Mylvaganam S, DiCiccio JE, Trimble WS, Bassik MC, Bonifacino JS et al. 2019. Phagolysosome resolution requires contacts with the endoplasmic reticulum and phosphatidylinositol- 4-phosphate signalling. Nat Cell Biol 21: 1234–1247.

Martinez J, Cunha LD, Park S, Yang M, Lu Q, Orchard R, Li QZ, Yan M, Janke L, Guy C et al. 2016. Noncanonical autophagy inhibits the autoinflammatory, lupus-like response to dying cells. Nature 533: 115–119.

McCray BA, Skordalakes E, Taylor JP. 2010. Disease mutations in Rab7 result in unregulated nucleotide exchange and inappropriate activation. Hum Mol Genet 19: 1033–1047.

Montano F, Grinstein S, Levin R. 2018. Quantitative Phagocytosis Assays in Primary and Cultured Macrophages. Methods Mol Biol 1784: 151–163.

Morris C, Foster OK, Handa S, Peloza K, Voss L, Somhegyi H, Jian Y, Vo MV, Harp M, Rambo FM et al. 2018. Function and regulation of the Caenorhabditis elegans Rab32 family member GLO-1 in lysosome-related organelle biogenesis. PLoS Genet 14: e1007772.

Nakae I, Fujino T, Kobayashi T, Sasaki A, Kikko Y, Fukuyama M, Gengyo-Ando K, Mitani S, Kontani K, Katada T. 2010. The arf-like GTPase Arl8 mediates delivery of endocytosed macromolecules to lysosomes in Caenorhabditis elegans. Mol Biol Cell 21: 2434–2442.

Nguyen JA, Yates RM. 2021. Better Together: Current Insights Into Phagosome-Lysosome Fusion. Front Immunol 12: 636078.

Niwa S, Lipton DM, Morikawa M, Zhao C, Hirokawa N, Lu H, Shen K. 2016. Autoinhibition of a Neuronal Kinesin UNC-104/KIF1A Regulates the Size and Density of Synapses. Cell Rep 16: 2129–2141.

Niwa S, Tao L, Lu SY, Liew GM, Feng W, Nachury MV, Shen K. 2017. BORC Regulates the Axonal Transport of Synaptic Vesicle Precursors by Activating ARL-8. Curr Biol 27: 2569–2578 e2564.

Pu J, Keren-Kaplan T, Bonifacino JS. 2017. A Ragulator-BORC interaction controls lysosome positioning in response to amino acid availability. J Cell Biol 216: 4183–4197.

Pu J, Schindler C, Jia R, Jarnik M, Backlund P, Bonifacino JS. 2015. BORC, a multisubunit complex that regulates lysosome positioning. Dev Cell 33: 176–188.

Qi W, Yan Y, Pfeifer D, Donner VGE, Wang Y, Maier W, Baumeister R. 2017. C. elegans DAF-16/FOXO interacts with TGF-ss/BMP signaling to induce germline tumor formation via mTORC1 activation. PLoS Genet 13: e1006801.

Rosa-Ferreira C, Munro S. 2011. Arl8 and SKIP act together to link lysosomes to kinesin-1. Dev Cell 21: 1171–1178.

Sancak Y, Bar-Peled L, Zoncu R, Markhard AL, Nada S, Sabatini DM. 2010. Ragulator-Rag complex targets mTORC1 to the lysosomal surface and is necessary for its activation by amino acids. Cell 141: 290–303.

Sasaki A, Nakae I, Nagasawa M, Hashimoto K, Abe F, Saito K, Fukuyama M, Gengyo-Ando K, Mitani S, Katada T et al. 2013. Arl8/ARL-8 functions in apoptotic cell removal by mediating phagolysosome formation in Caenorhabditis elegans. Mol Biol Cell 24: 1584–1592.

Saxton RA, Sabatini DM. 2017. mTOR Signaling in Growth, Metabolism, and Disease. Cell 169: 361–371.

Stenmark H, Parton RG, Steele-Mortimer O, Lutcke A, Gruenberg J, Zerial M. 1994. Inhibition of rab5 GTPase activity stimulates membrane fusion in endocytosis. EMBO J 13: 1287–1296.

Yang HY, Mains PE, McNally FJ. 2005. Kinesin-1 mediates translocation of the meiotic spindle to the oocyte cortex through KCA-1, a novel cargo adapter. J Cell Biol 169: 447–457.

Yu L, McPhee CK, Zheng L, Mardones GA, Rong Y, Peng J, Mi N, Zhao Y, Liu Z, Wan F et al. 2010. Termination of autophagy and reformation of lysosomes regulated by mTOR. Nature 465: 942–946.

